# Landscape of reversion alterations in homologous recombination genes reveals evolutionary constraints

**DOI:** 10.64898/2026.05.19.726319

**Authors:** Brennan Decker, Saumya D Sisoudiya, James Thornton, Chenming Cui, Taku Harada, Shahlo O Solieva, Russell W Madison, Arindam Datta, Tao Shi, Fabio Giuntini, Zoe Fleischmann, Mark Hartman, Shannon T Bailey, Jeffrey A Leibowitz, Ethan S Sokol, Alexa B Schrock, Garrett M Frampton, Fergus J Couch, Kim A Reiss, Susan M Domchek, Gregory R Bowman, Roger A Greenberg, Smruthy Sivakumar, Katherine L Nathanson

## Abstract

Reversion mutations (REVs) restore homologous recombination repair (HRR) and confer resistance to PARP inhibitors (PARPi) in HR-deficient cancers. Yet, their prevalence, mechanisms, and biological constraints remain undefined. We analyzed genomic profiling of 609,464 tissue and liquid biopsy samples across multiple cancer types to delineate the pan-cancer landscape of REVs. REVs were identified in eight HRR genes, most frequently *BRCA2* and *BRCA1* and notably never in *ATM* or *CHEK2*. REVs exclusively impacted truncating pathogenic variants, predominantly through large in-frame and exon-level deletions, associated with repetitive sequences. Conserved functional domains are relatively depleted of REVs. Structural modeling and functional studies support that exon-level deletions preserve critical domain architecture and confer PARPi resistance. The study establishes HRR reversion as a structurally permissive, yet evolutionarily constrained, resistance mechanism with implications on response, monitoring, and therapeutic strategy.

**One Sentence Summary:** Homologous recombination reversion evolves through structurally tolerated, microhomology-driven alterations that restore DNA repair under therapeutic selection across cancers.

## INTRODUCTION

Pathogenic variants (PV) in genes encoding components of the homologous recombination repair (HRR) pathway confer increased risk for multiple malignancies, most prominently breast and ovarian cancer (*1*). Tumors arising in PV carriers typically exhibit biallelic HRR gene inactivation, resulting in loss of high-fidelity, sister-chromatid-guided repair of double-stranded DNA breaks (*2*). This homologous recombination deficiency (HRD) creates a therapeutic vulnerability to agents that generate lesions that require HR repair to complete DNA replication, including platinum-based chemotherapy and poly (ADP-ribose) polymerase inhibitors (PARPi) (*3, 4*). PARPi have demonstrated substantial clinical benefit across frontline, maintenance, recurrent, and adjuvant settings in HRD-associated breast, ovarian, prostate, and pancreatic cancers with germline or somatic *BRCA1* or *BRCA2* PVs, and to a lesser extent PVs in *PALB2*, *RAD51C*, and *RAD51D* (*5–8*). These data have led to multiple FDA approvals, establishing PARPi as a core therapeutic strategy in these malignancies (*7, 9–12*).

Despite initial sensitivity to platinum chemotherapy and PARPi, resistance almost invariably emerges (*13*). The most frequently observed mechanism of acquired resistance is development of a reversion mutation (REV), in which a secondary genetic alteration reverses the effect of a PV in the affected HRR gene, thereby partially or fully reconstituting HRR and mediating partial or complete therapy resistance (*13–19*). The evolutionary pressure that produces REVs favors restored HRR function rather than restoration of the wild-type protein and does not necessarily require preservation of canonical gene architecture (*16*), suggesting that REVs may be permissive of substantial structural disruption if they remain compatible with sufficient, even if attenuated, HRR activity.

Although REVs are widely recognized as central mediators of therapeutic resistance, reported frequencies following platinum or PARPi exposure vary substantially, ranging from approximately 10% to 40% across studies (*13, 16, 17, 20–24*). Most analyses have been limited by small cohorts, aggregation across heterogeneous patient populations or assays, restriction to individual tumor types or genes, and inconsistent incorporation of LBx data. Limited investigation of REVs in LBx has constrained detection of polyclonal and temporally evolving resistance events dynamically arising under therapeutic selection. Emerging evidence suggests gene- and allele-specific patterns of reversion, including higher REV rates in *BRCA2* than *BRCA1*, enrichment of large deletions within central exons, and relative sparing of highly conserved C-terminal domains (*16, 25*). Whether such domain-level constraints and structural tolerances generalize across tumor lineages remains unexplored. Even within *BRCA1* and *BRCA2*, clinical data remain sparse, limiting understanding of REV prevalence, structural diversity, and domain-level constraints. Notably, large *BRCA2* exonic deletions have been identified as frequent resistance mechanisms in preclinical models (*19*) but have not been identified in patient-derived cohorts. Larger, clinically annotated datasets are therefore required to define the pan-cancer landscape of HRR REVs across genes, tumor types, and pathogenic variant classes.

Herein, we analyze comprehensive genomic profiling (CGP) data from a large pan-cancer cohort of tumor tissue (TBx) and liquid biopsy (LBx) samples to define the prevalence, distribution, and structural features of REVs. We identify HRR genes in which REVs are detected or conspicuously absent and quantify REV rates by tumor type, gene, pathogenic variant (PV) class, and allele. The breadth of observed PV-REV pairs enable resolution of gene-, domain-, and mutation-specific resistance patterns. Finally, we identify numerous REVs mediated by exon or multi-exon deletions and use AI-enabled structural modeling to demonstrate that selected *BRCA1* and *BRCA2* deletions preserve or restore critical functional domain architecture, consistent with restored function and therapy resistance, with experimental validation for *BRCA2* exon 11 deletions.

## RESULTS

### Spectrum and prevalence of HRR reversion mutations

This cross-sectional study analyzed comprehensive genomic profiling data from 609,464 unique patients, including 516,279 tissue biopsies (TBx) and 93,185 liquid biopsies (LBx) (**fig. S1**). Pathogenic variants (PV) were identified in fourteen HRR genes associated with FDA-approved PARPi companion diagnostic claims in 101,674 patients (16.7%), including 75,417 TBx (14.6%) and 26,257 LBx (28.2%). TBx PV rates ranged from 0.22% for *CHEK1* to 4.1% for *ATM*; higher LBx prevalences were largely driven by clonal hematopoiesis in *ATM* and *CHEK2* (**Fig. 1A; table S1**).

**Fig. 1.**
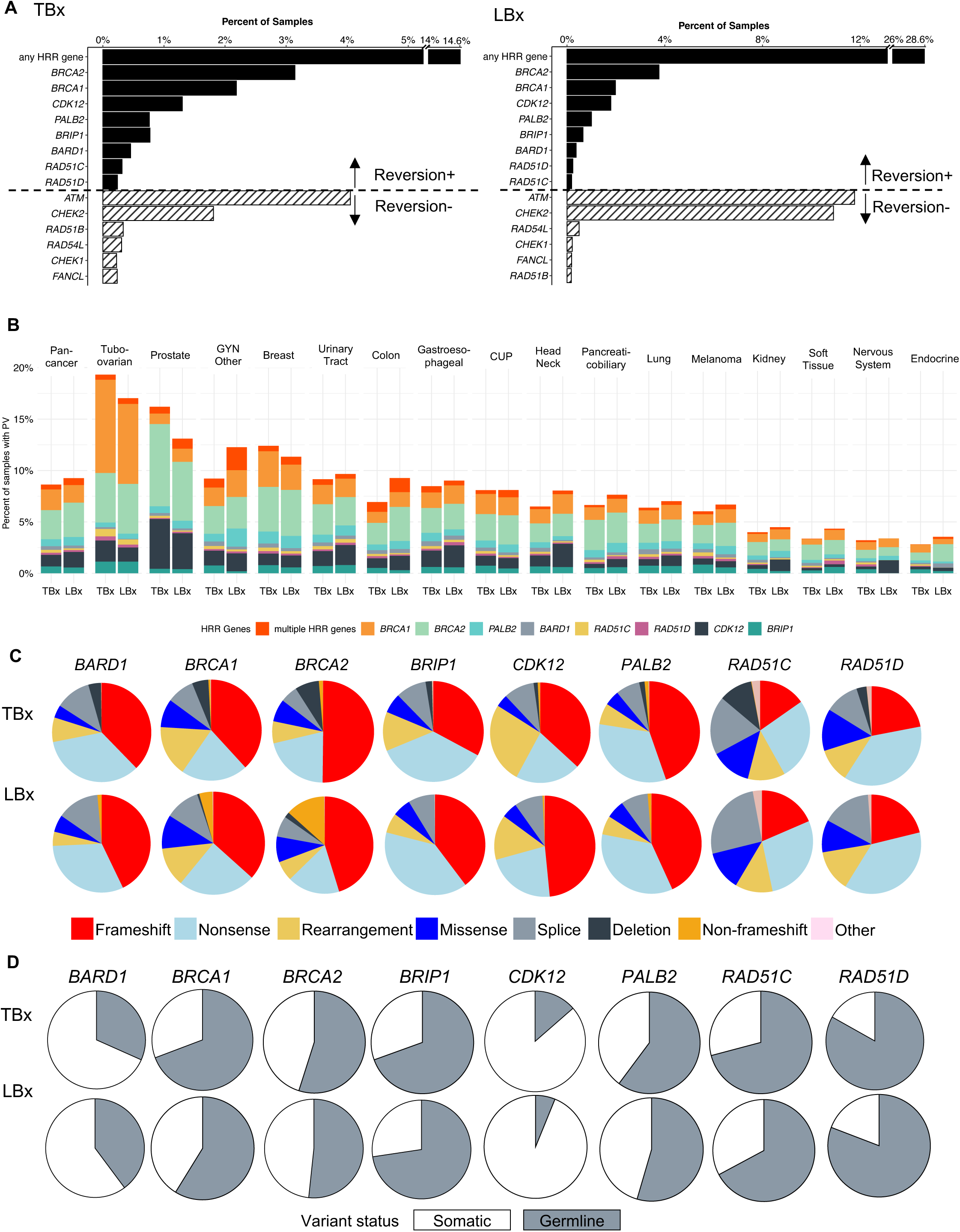
Landscape of pathogenic variants (PVs) for genes in which reversion mutations (REVs) are observed for tissue biopsy (TBx) and liquid biopsy (LBx) specimens. **A)** Pan-cancer, sample-level PV rates for 14 genes included in FDA-approved PARPi companion diagnostic (CDx) indications. Genes are grouped by whether REVs were observed at least once in that gene. REVs were exclusively observed in eight genes, referred to as REV8. **B)** PV rates by cancer type for the REV8 genes. Samples with high TMB were removed, as they frequently harbor monoallelic passenger mutations that do not predict functional readouts of HRD or response to PARPi therapy (*61*) see methods and Figure S1). No REV+ samples were removed. **C)** Breakdown of PV type by gene. **D)** Predicted germline versus somatic origin of PVs by gene for the subset of samples for which a confident result was obtained.

Reversion mutations (REVs) were detected in eight genes: *BRCA1, BRCA2, PALB2, RAD51C, RAD51D, BARD1, BRIP1,* and *CDK12* (hereafter “REV8”). *BRCA2* (TBx 3.1%; LBx 3.8%) and *BRCA1* (TBx 2.2%; LBx 2.0%) harbored the most frequent PV. Only samples with ≥1 PV in a REV8 gene were considered at risk for reversion. Among 52,853 such samples, 701 (1.3%) were REV-positive (REV+), including 458 TBx and 243 LBx (**table S2**). Patients with REV+ tumors were significantly younger and more likely female (**table S2**), consistent with an integral role for platinum and PARPi therapies in clinical management of breast and ovarian cancer.

### Distribution of REV-eligible PV across cancer types, variant classes, and ancestry

PV in REV8 genes were most prevalent in tubo-ovarian cancers (TBx 19.5%) and exceeded 10% of TBx and LBx samples in prostate, breast, and other gynecologic malignancies (**Fig. 1B; table S3**). PV spanned all major variant classes, including truncation, rearrangement, large deletion, missense, and in-frame indels, with similar distributions between TBx and LBx (**Fig. 1C; table S4**). *RAD51C* and *RAD51D* exhibited relatively fewer frameshift PVs compared with other genes.

Germline versus somatic origin was predicted for 41.5% of PVs (25,583/61,566), revealing predominantly somatic alterations in *CDK12* and *BARD1*, whereas germline PV predominated in other genes (range 55% [TBx] in *BRCA2* to 83.2% [TBx] in *RAD51D*; **Fig. 1D; table S5**). REV8 PV frequencies were comparable across genomic ancestry superpopulations (**Fig. S3; table S6**).

### Pan-cancer landscape of REV frequency by biopsy type, cancer type, and gene

REV rates were significantly higher in LBx than TBx (2.9% vs 1.0%) and highest in LBx with tumor fraction >1% (3.8%; P<0.0001), reflecting enrichment for advanced-stage disease (P<0.0001; **Fig. 2A; table S7**). TBx samples exhibited a median of one REV per sample (range 1-4), with 91.5% harboring a single REV, whereas LBx samples had a median of two REVs (range 1-39), and 53.5% contained multiple independent REVs (**Fig. 2B**). One LBx harbored 39 distinct in-frame deletion REVs (range 2-1,179 amino acids [AA] in length) correcting a single PV (**fig. S4A**).

**Fig. 2.**
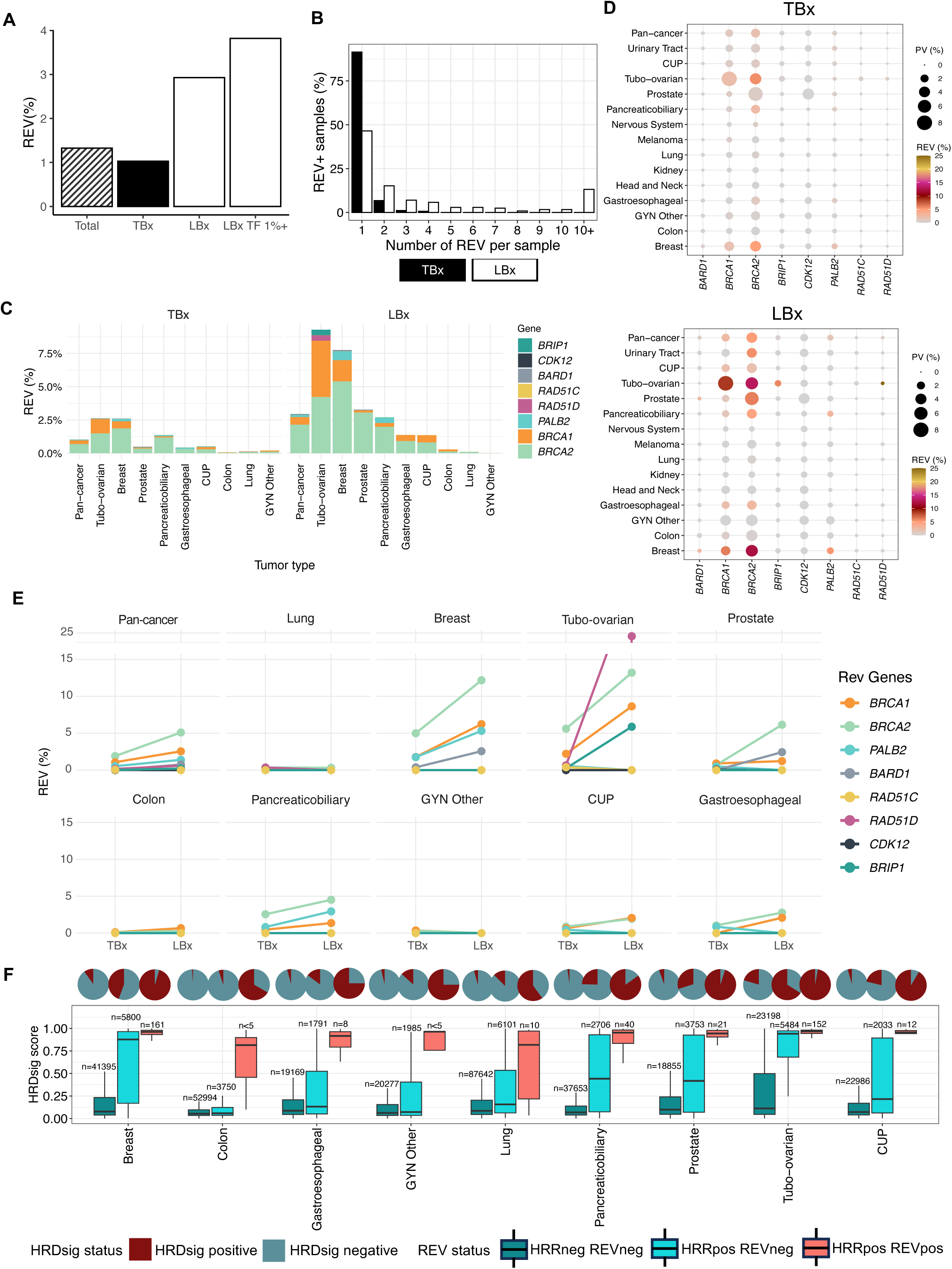
Pan-cancer landscape of REVs for TBx and LBx samples. **A)** REV rate, calculated as the number of REV+ samples divided by the total number subjects with REV8 PV. **B)** REV rate by cancer type, with color representing the gene harboring a REV. **C)** TBx vs LBx REV rates, stratified by cancer type and gene. **D)** Prevalence of PVs in REV8 genes (circle size) and gene-specific REV rate (circle color) by cancer type. **E)** Number of REVs observed per sample, stratified by sample matrix. **F)** HRDsig score, a machine learning-based measure of HRD-associated copy number features, for cancer type of interest, stratified by REV8 gene PV status and REV status. Pie charts show the overall HRDsig status, with scores >0.7 validated as positive, whereas boxplots depict the distribution of scores. HRRneg / REVneg = Samples with no HRR gene PVs and no REV; HRRpos / REVneg = samples with one or more PVs in one or more REV8 genes, but no REV detected; HRRpos / REVpos = samples with one or more PVs in one or more REV8 genes and one or more REVs detected in the same gene.

REV rates were highest in tubo-ovarian (TBx 2.6%; LBx 9.1%) and breast cancers (TBx 2.5%; LBx 7.7%), followed by pancreaticobiliary and prostate malignancies (**Fig. 2C; table S8**). REVs were more rarely detected in a long tail of other malignancies. *BRCA2* exhibited the highest gene-specific REV rate (TBx 1.9%; LBx 5.1%), followed by *BRCA1* and *PALB2*; REVs in the other genes were rare (**Fig. 2D; table S9**). Gene-specific REV rates further differed by cancer type. Although *BRCA1* PV were more prevalent in tubo-ovarian cancers, *BRCA2* PV were significantly more likely to revert (5.6% vs 2.2%; P<0.0001), a pattern consistently observed across tumor types. Gene-specific REV rates were universally higher in LBx than TBx (**Fig. 2E; table S10**).

Across all tumor groups, REV+ samples exhibited significantly higher HRDsig scores (a machine learning-based quantitative assessment of copy number patterns associated with HRD (*26*)) than REV-negative samples (P<0.0001; **Fig. 2F; table S11**). In malignancies less commonly associated with HRD, HRDsig scores varied more widely among REV8+ than REV+ samples, consistent with enrichment for monoallelic passenger PVs.

### Pathogenic variant class dictates REV rate and restoration mechanisms

Although all classes of PVs were detected in REV8 genes, REVs exclusively corrected truncating PVs (frameshift, nonsense, or splice variants). Frameshift PVs exhibited the highest REV rates, exceeding nonsense PVs by 1.5-fold and splice PVs by approximately 8-fold across both TBx and LBx (all P<0.0001). REVs were not observed for missense, deletion, rearrangement, or non-frameshift PVs (**Fig. 3A; table S12**). The total number of reverted PVs (n=725) exceeded the number of REV+ samples as 18 samples harbored two or more independently reverted PVs within the same gene (example in **fig. S4B**).

**Fig. 3.**
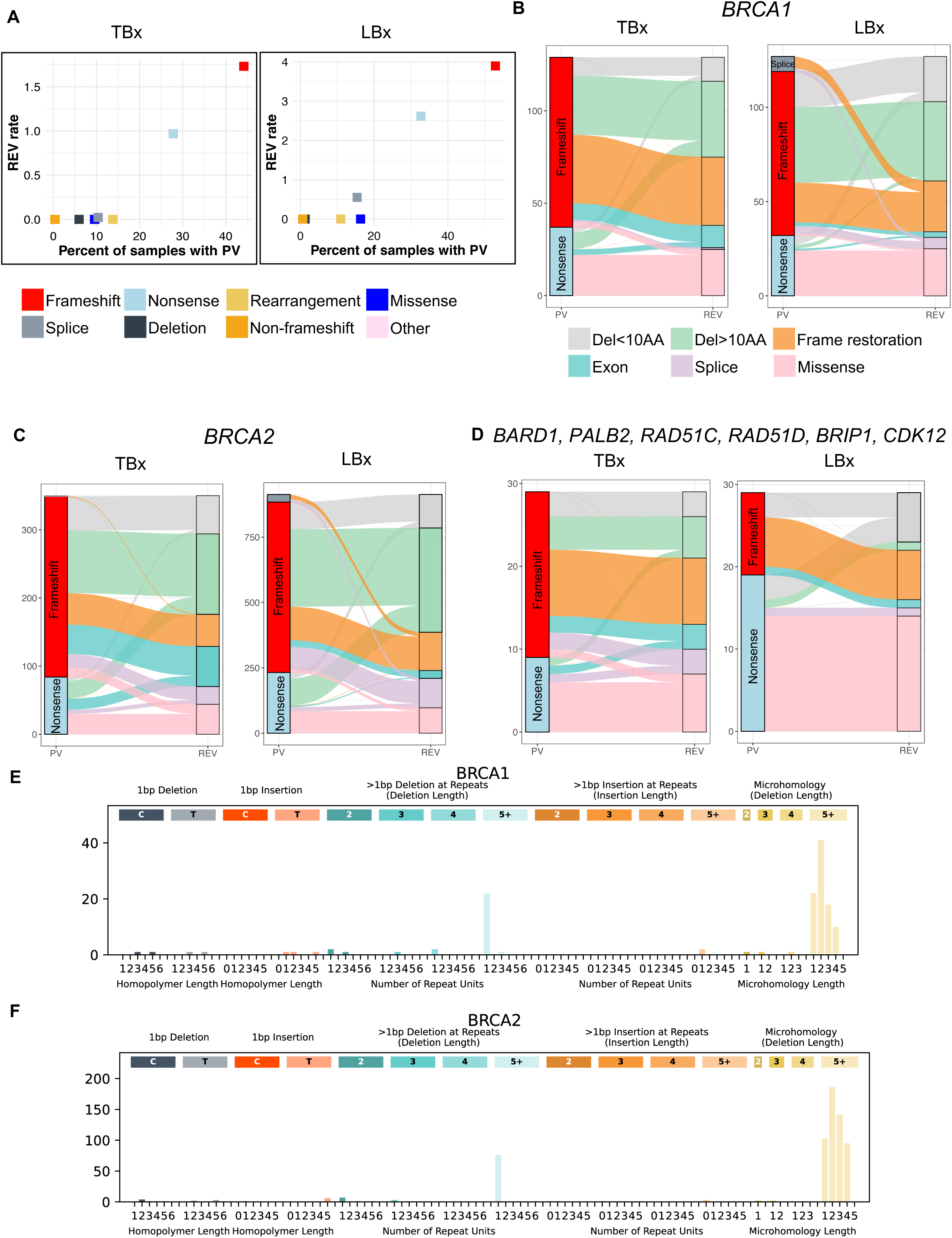
REV rates and mechanisms by PV type. **A)** PV prevalence versus REV rate by predicted effect of PV. **B-D)** Frequency of reversion mechanisms for all observed REVs, broken down by PV type for *BRCA1* (**B**), *BRCA2* (**C**), and the other REV8 genes (**D**). **E-F)** Indel signatures for mutations causing REV in *BRCA1* (**E**) and *BRCA2* (**F**), including counts of mutations in three categories: 1-bp insertion or deletion in the context of increasing homopolymer length (slate and fire), insertions or deletions larger than 1-bp in repetitive regions by length of the repeat (light blue/orange), and deletions that occur with microhomology by length of the microhomology region (yellow). Complex deletion-insertion REV were excluded.

Among 1,575 total REVs (**Fig. 3B–D; table S12**), in-frame deletions, including exon-level deletions, were the most common mechanism (60%; n=944). Exon-level deletions accounted for 7% (n=108) of REVs overall, with lower detection rates for this mechanism in LBx (2%; n=34), presumably due to challenges in detecting this REV type when ctDNA shed is low. Sub-exon-level deletion lengths ranged from 1-1,221 amino acids, with 72% (n=606/836) ≥10 amino acids and a strong bias toward correction of frameshift PVs (P<0.0001). In contrast, missense substitutions predominated among REVs correcting nonsense PVs (43% vs 2.2% for frameshift; P<0.0001). REVs correcting splice PVs were rare and mechanistically diverse with some restoring wild-type splice junctions and others introducing novel, frame correcting predicted splice donors or acceptors (**Fig. S2**).

REV mechanism usage varied by gene. In-frame deletions accounted for 62.6% of *BRCA2* REVs (n=790/1261) but only 32.8% (n=19/58) across *PALB2*, *RAD51C*, *RAD51D, BARD1, BRIP1,* and *CDK12* (P<0.0001). Deletion length was significantly greater in *BRCA2* than *BRCA1* (medians of 81 vs 21 AAs; P<0.0001; **Fig. S5A**). In both genes, long deletions were strongly enriched in the large central exons and rarely observed elsewhere. In *BRCA2*, 90% of inframe deletion REVs were localized to exon 11 and were significantly longer than those elsewhere in the gene (**fig. S5B**, median of 93 AA vs 11 AA, respectively; P<0.0001). Deletion REVs in *BRCA1/2* were frequently mediated by microhomology, particularly within large central exons (**Fig. 3E, F**), whereas homopolymers and other repeats predominated elsewhere (**fig. S5C, D**). Notably, the BRCA2 exon 11 deleted allele was expressed at considerably higher levels than full length BRCA2 (**Fig 4E; fig S9C**), which may explain how restored PARPi resistance can occur despite complete loss of the BRCA2 BRC repeats.

**Fig. 4.**
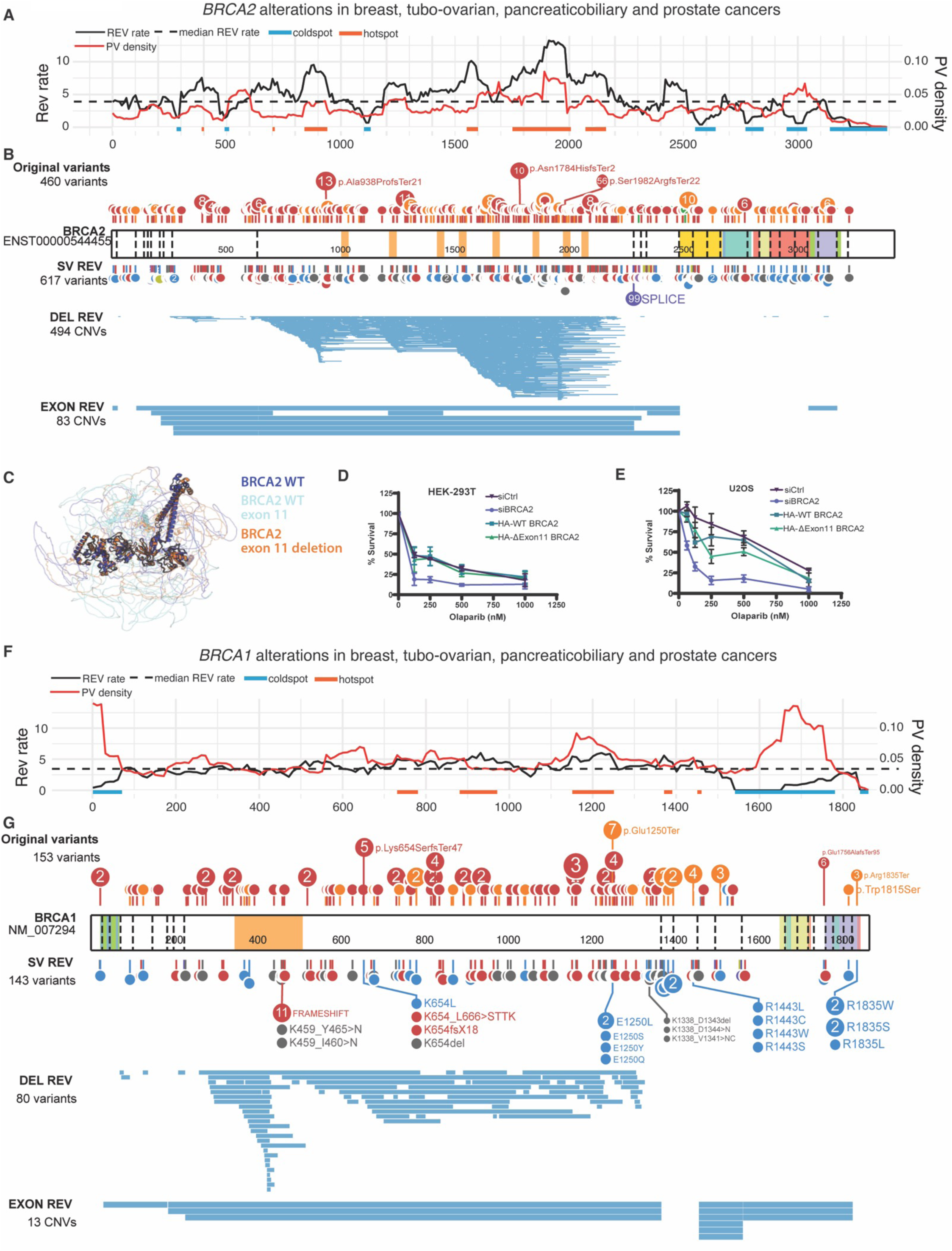
Relationships between PVs and REVs in *BRCA2* and *BRCA1*. **A)** *BRCA2* PV density and REV rate within a 100-AA sliding window by PV AA position for breast, tubo-ovarian, prostate, and pancreatic cancers. **B)** Reverted *BRCA2* PV type and position (track above the gene model) and REV mechanism and position (tracks below the gene model), with the upper track containing REV <10 AA in length, the middle track depicting inframe deletion REVs within exons and spanning >10 AA in length, and the bottom track displaying exon-level or larger deletion REVs. **C)** Comparison of predicted structure for WT BRCA2 (dark blue with exon 11 in light blue) and BRCA2 without exon 11 (orange). Results of colony forming assay comparing HA-WT BRCA2 and HA-ΔExon11 BRCA2 in **D)** U2OS and **E)** 293T cells with siBRCA2 knockdown. **F)** *BRCA1* PV density and REV rate within a 100-AA sliding window by PV AA position for breast, tubo-ovarian, prostate, and pancreatic cancers. **G)** Reverted *BRCA1* PV type and position (track above the gene model) and REV mechanism and position (tracks below the gene model), with the upper track containing REV <10 AA in length, the middle track depicting inframe deletion REVs within exons and spanning >10 AA in length, and the bottom track displaying exon-level or larger deletion REVs.

### Regional REV constraints, structural tolerance, and functional validation

REV rates varied markedly across gene regions. In *BRCA2*, REV rate peaked within exon 11, exceeding 8% between codons 1781-2101, with rates below 2% and fewer long deletions in the 3′ region encoding essential functional domains (six after exon 11 vs 571 in exons 1-11) (**Fig. 4A, B, table S13**). Two cold spots were observed in exon 10, at nucleotides 291-391, containing the N terminal DNA binding domain (*27*) and 491-601, a region without delineated function. Given the prevalence of large exon-removing REVs, including *BRCA2* REVs predicted to eliminate exon 11 via deletion (5%; n=63/1202) or splice site interruption (8.4%; n=101/1202), AlphaFold modeling was used to assess their structural impact. Models for deletion constructs predicted preserved spatial configuration of BRCA2 catalytic domains (exons 15-25), with minimal differences in Cα RMSD (0.94-2.97Å) and pLDDT scores (**Fig. 4C; fig. S6-8**). Consistent with these predictions, transient expression of exon 11-deleted BRCA2 conferred PARPi resistance comparable to full-length BRCA2 in colony formation assays in U2OS and 293T cells with BRCA2 knockdown (**Fig. 4D, E; fig. S9**). Exon 11-deleted BRCA2 also co-localized with RAD51 foci after treatment with PARPi (**fig. S10**).

In *BRCA1*, REV rates were low in the 5′ region (1-161 AA) (**Fig. 4F, table S14**), consistent with reports of translation reinitiation at AUG297 (*28, 29*); resultant peptides lack the RING domain but do not appear to require a REV to restore partial protein function. REVs were depleted in the 3’ (1,541 AA and beyond) of the gene, including the BRCT-containing region, despite enrichment for PVs in this region (**Fig. 4F, table S14**). Exon 10 harbored recurrent in-frame deletion REVs with a shared termination point (n=465 AA) near the nuclear localization sequence, which may represent homology-mediated recurrent breakpoints. BRCA1 AlphaFold structural modeling of exon-level deletions revealed minimal differences in the configuration of the critical exons 2-5 (0.28-0.68 Å) and 16-23 (0.14-0.36 Å) by both Cα RMSD and pLDDT (**fig. S11-13**). *PALB2* exhibited lower REV rates mid-protein and in the WD40 domain, although interpretation was limited by sample size (**fig. S14**).

### Reversion frequency of recurrent pathogenic variants

REV rates varied widely across individual recurrent PVs (range 0%-27.6%; **table S15**). *BRCA2* p.Gln397LeufsTer25 (n=8/29; 27.6%) and p.Thr2766AsnfsTer11 (n=6/29; 21%) exhibited the highest REV rates (**table S15**). The Ashkenazi Jewish (AJ) founder allele *BRCA2* p.Ser1982ArgfsTer22 had a 14% (n=56/402) REV rate, with 190 total (157 unique) REVs, of which 33 were recurrent across samples (**fig. S4C**), raising the possibility of shared genomic mechanisms for developing REV. In contrast, the AJ founder allele *BRCA1* p.Glu23ValfsTer17 was the most frequently observed PV in the gene (n=385) and exhibited a REV rate of only 0.5%. Common missense PVs, such as *BRCA1* p.Cys61Gly, *BRCA2* p.Asp2723His and p.Arg2842Cys, were never reverted (**table S15**).

## DISCUSSION

REVs represent a dominant mechanism of acquired resistance to platinum-based chemotherapy and PARPi in HRD cancers, yet their biological constraints, structural requirements, and prevalence across tumor types have remained incompletely defined (*14, 18*). By evaluating PVs and REVs with a single consistent methodology via comprehensive genomic profiling of more than 600,000 TBx and LBx samples across cancer types, this study defines the pan-cancer landscape of HRR REVs at unprecedented scale and granularity. REV rates in this study are substantially higher than reported in other pan-cancer analyses (*30, 31*), likely due to the inclusion of data from liquid biopsies and systematic identification of large exonic deletions. REVs were identified more frequently in LBx than TBx, with the highest rates in LBx with tumor fraction over 1%, likely due to sampling at later stages upon progression on targeted therapy. REVs were most commonly observed in tubo-ovarian and breast cancer samples, most frequently in *BRCA2*, followed by *BRCA1*, with very low rates in the other genes. These data establish reversion as a structurally permissive, evolutionarily constrained, and dynamically evolving resistance process governed by domain architecture, error-prone DNA repair, and allele-specific context.

The restricted distribution of reversion mutations across HRR genes reinforces the link between HRD-driven therapeutic sensitivity and reversion-mediated resistance, with detection or absence of REVs potentially serving as a potential indicator of therapeutic susceptibility or alternative mechanisms of resistance. Despite PVs being detected across fourteen DNA repair genes associated with FDA approvals, REVs were confined to eight genes. Notably, REVs were absent in *ATM* and *CHEK2*, genes not associated with HRD genomic signatures (*32*) and for which evidence for PARPi benefit has been inconsistent (*33–35*). These findings support the interpretation that pathogenic variants in these genes do not generate the selective pressures required for reversion in response to PARPi or platinum exposure.

PVs had non-equivalent probabilities of REV, with rates stratified by variant type, variant position within the gene, and even specific alleles, which can be quantified. Across all genes, REVs were observed exclusively for truncating PVs (frameshift, nonsense, and splice alteration) and were never detected for pathogenic missense, inframe indels, or genomic rearrangements. This absence is unlikely to reflect technical limitations or sample size, given frequent detection of REVs affecting nonsense or frameshift PVs and numerous observations of established PVs in these classes. For missense PVs, lack of reversion may reflect fundamental biological constraints, since they are in highly conserved domains in which secondary mutations may be structurally or functionally intolerable. Alternatively, missense PV alleles may retain sufficient residual function to reduce selective pressure for reversion. Experimental models demonstrating hypomorphic behavior of *BRCA1* missense variants and alternative resistance mechanisms support this interpretation (*36–38*). The observed differences in REV frequency across genes and PV types, especially for specific variants, raises the possibility of stratifying patients by reversion-susceptible versus reversion-resistant PVs to refine patient selection, improve interpretation of resistance mechanisms, and inform trial design for therapies targeting DNA damage response pathways.

A central finding is that HRR functional restoration through REV does not require retention or reconstruction of the entirety of the canonical gene architecture. Instead, resistance is frequently achieved through large in-frame and exon-level deletions that preserve essential catalytic domains while simultaneously eliminating substantial portions of the protein, which can be inclusive of some functional domains. This phenomenon is most striking in *BRCA2*, in which deletion or splice-mediated skipping of exon 11, removing all BRC repeats, was the most common REV mechanism (13.4% of total REVs). Structural modeling demonstrated preservation of the C-terminal domains required for BRCA2 activity despite extensive central deletions, and functional assays confirmed that BRCA2 Δexon 11 confers PARPi resistance comparable to full-length protein, confirming prior preclinical observations that large intragenic deletions can restore function leading to therapy resistance (*19, 29, 39*). Brca2 floxed Δexon 11 mouse models have been used to model BRCA2 loss in multiple studies (*40, 41*); these data suggest that these models may be functionally hypomorphic rather than null, potentially providing an explanation for therapeutic resistance seen in these models.

REV patterns across genes further reveal strong domain-level constraints on resistance evolution. Regions enriched for repetitive elements, particularly the large central exons of *BRCA1* and *BRCA2*, tolerated extensive deletions and exhibited high REV rates, whereas conserved domains were markedly depleted of REV despite substantial PV burden. These cold spots suggest that loss of these critical functional domains imposes prohibitive fitness costs, limiting the evolutionary paths available for resistance. Such domain-level constraints are consistent with prior structural and functional studies demonstrating the essentiality of BRCA2 C-terminal domains for RAD51 binding, DNA binding, and replication fork protection (*42, 43*) and NTD domain at the N-terminus (*27*). Multiple studies have demonstrated that variants in *BRCA1* and *BRCA2* are associated with a range of cancer risk (*44–46*), with some missense and splice variants associated with lower risk (*47*). Together, these findings raise the possibility that some alterations may cause sufficient loss of HRR function to promote development of cancer while simultaneously retaining adequate residual function to evade synthetic lethality (**Fig. 5**).

**Fig. 5.**
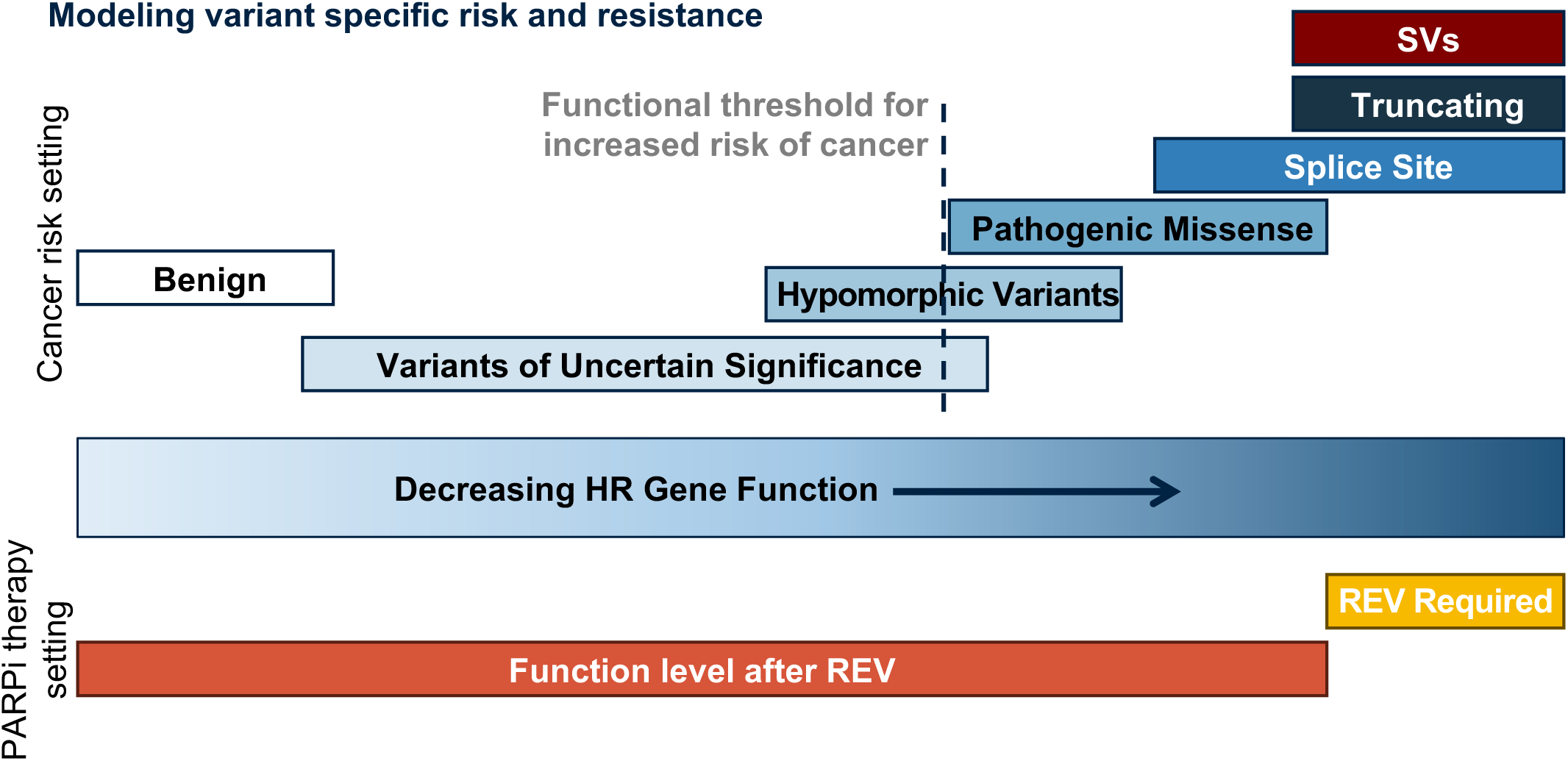
Model for variant specific risk and resistance. Emerging data on pathogenic variants in BRCA1 and BRCA2 suggests that they confer a spectrum of loss of function and level of cancer susceptibility risk, predicted by the mutation type and its location within the gene. In parallel, the data herein suggests that although some variant types, generally with lower cancer risk, have a decrement in protein function sufficient for cancer susceptibility, they retain enough DNA damage repair function that reversion variants do not emerge as mechanisms of therapeutic resistance.

The mutational mechanisms underlying reversion support a model in which resistance emerges as a direct consequence of HRR deficiency itself. Deletion-based reversion events were frequently mediated by long stretches of microhomology, consistent with reliance on microhomology-mediated end joining and related error-prone repair pathways that become upregulated in the absence of HRR. This mechanism is consistent with HRD-associated indel signatures and prior observations linking microhomology-mediated repair to REV (*48*). For this mechanism, REVs represent an emergent outcome of sustained therapeutic pressure acting on HR-deficient genomes rather than an independent resistance pathway.

Analysis of LBx samples highlights REVs as a dynamic, polyclonal evolutionary process. REV rates were higher in LBx than TBx, particularly in samples with higher tumor fraction, reflecting enrichment for advanced disease and prolonged therapy exposure. Many LBx samples harbored multiple independent REVs, and in extreme cases dozens of REVs were detected in a single sample. These findings extend prior cfDNA-based studies showing that resistance-associated REVs can be detected longitudinally and often emerge in parallel under therapeutic selection (*17, 49*).

This study has limitations. Its cross-sectional design limits temporal inference regarding the sequence of exposure to therapy and emergence of resistance events, and functional validation was necessarily restricted to a subset of observed mechanisms. REVs in less frequently altered HRR genes were rare, constraining gene-specific analyses. Patients in some cancer types and with driver PVs in some HR genes are unlikely to be treated with platinum or PARPi making observed REV rates difficult to interpret in these settings. Future longitudinal studies integrating serial liquid biopsy, expanded functional interrogation across HRR genes, and detailed treatment histories will be essential to fully resolve the dynamics of resistance evolution.

In summary, this pan-cancer analysis reveals REVs as a structurally permissive, yet evolutionarily constrained mechanism of therapeutic resistance. By defining the architectural, mechanistic, and allele-specific principles governing REVs across genes, sample matrices, and cancer types, this work provides a framework for anticipating resistance, interpreting emerging resistance variants, and improving the durability of HRD-directed therapies.

## MATERIALS AND METHODS

### Study design and samples

Approval for this study, including a waiver of informed consent and a HIPAA waiver of authorization, was obtained from the Western Institutional Review Board (Protocol No. 20152817). Patients with comprehensive genomic profiling (CGP) by Foundation Medicine, Inc. (Boston, MA, USA) ordered during routine clinical care at institutions across the United States from 2012 to 2024 were included (**fig. S1**). Tumor type was determined by a board-eligible pathologist based on integration of available information including pathology report information, submitting physician diagnosis, immunohistochemistry, and genomic findings. Basic demographic information (age, sex) and sample information (sample site, date of sample collection) were extracted from test requisitions and pathology reports. When a patient had more than one CGP test by Foundation Medicine, the sample that passed all quality control (QC) metrics, or the first reversion-positive sample if all samples passed QC metrics, was chosen for primary analysis. Patients with multiple samples with the same diagnosis, identified by evaluation of genotypes at known polymorphic sites, were compared across time points.

### Comprehensive genomic profiling

CGP was performed in a CLIA-certified, CAP-accredited laboratory (Foundation Medicine, Cambridge, MA, USA) by next generation sequencing using adaptor-ligation and hybrid capture, as previously described (*50, 51*). For patients with solid tumor TBx, sequencing was run on DNA extracted from formalin-fixed paraffin-embedded (FFPE) samples and 187-416 genes were evaluated for all classes of alterations (short variants such as single nucleotide variants and short insertions/deletions, copy number alterations such as gene amplifications and homozygous deletions, and large genomic rearrangements). For patients with LBx, sequencing was performed on DNA extracted from peripheral whole blood, and 62-324 genes were evaluated for all classes of genomic alterations (*52*).

Somatic/germline status for short variants was determined from tumor-only sequencing using the previously described somatic-germline-zygosity algorithm (*53*).

Microsatellite instability and tumor mutation burden were analyzed (*54, 55*). For both TBx and LBx, mutation signatures were determined for any case with at least 10 non-driver somatic mutations by analyzing the trinucleotide context using the Sanger COSMIC mutational signatures (*56*). Phenotypic HRD was assessed using HRDsig, a machine learning based algorithm designed to detect copy number features associated with HRD (**Supplemental Methods**) (*26*).

### Detection and classification of REV

To meet the definition of a REV, a variant must nullify the effect of the original PV. REVs were classified computationally using a bespoke research use only algorithm that analyzed two variants occurring within the same gene for potential interactions compatible with causing a REV. Samples with at least one pathogenic variant (PV) and at least two variants occurring within the same HR gene were identified and processed with the bespoke algorithm that tested for seven mechanisms of REV (**fig. S2**): 1) PV encompassed by an inframe deletion; 2) PV replaced by a missense substitution; 3) frameshift PV corrected by frame-restoring indel; 4) exon containing a PV predicted to undergo skipping due to splice site mutation; 5) PV removed by an exon-level deletion; 6) biallelic deletion of the region containing the PV; and 7) splice PV corrected by frame restoration or introduction of predicted novel splice site.

### Statistical methods and software

Comparison of age and tumor mutational burden (TMB) of patients with PVs, comparing those with and without REVs was performed using Wilcoxon rank sum test in Table 1. Comparison of categorical variables such as sex, ancestry, biopsy site, tumor stage, HRDsig status, and MSI status was performed using Pearson’s Chi-squared test or Fisher’s exact test, depending on number of samples across the different categories in Table 1. In-frame deletion REV lengths were compared using Wilcoxon rank sum test and Pearson’s Chi-squared test was used for all remaining statistical comparisons throughout the manuscript. Statistical significance was set at *P* ≤ 0.05.

### Structural modeling

Structure predictions were generated using ColabFold (*57, 58*) (https://github.com/YoshitakaMo/localcolabfold), which combines AlphaFold2 with MMseqs2 to accelerate prediction times. Multiple sequence alignments (MSAs) were computed on the MMseqs2 MSA server. The number of recycles was set to three, the default ColabFold value. We predicted five models per sequence and used Amber for structure relaxation. However, the following structures did not relax: BRCA2 wildtype, BRCA2 exons 24-25 deletion, BRCA2 exons 12-14 deletion, and BRCA2 exon 2 deletion. We calculated the average pLDDT (per-residue confidence metric) per exon by averaging the AlphaFold pLDDT scores across the residues within each exon. We report one standard deviation in the error bars. MDTraj (*59*) was used to index residues. We computed alpha carbon root-mean-square deviation (RMSD) values between the wildtype BRCA models and the deletion mutants by using the Molecular Software Library (MSL) (*60*). We compared the AlphaFold models with experimental BRCA1/2 structures, as detailed in **Supplementary Methods**.

### Functional studies

#### Transfection of Oligonucleotides

Wild type and mutant BRCA2 genes were synthesized then cloned into pcDNA3.1(+) plasmids by GenScript (Nanjing, China). All siRNA used in this study were synthesized by Horizon Discovery (Waterbeach, United Kingdom).

HEK293T cells were seeded at a density of 250,000 cells per well of a 6-well plate 24 hours before transfection. Oligonucleotides (siRNA and BRCA2 expression plasmids) were co-transfected into respective experimental groups using DharmaFECT Duo Transfection Reagent (Horizon Discovery T-2010-01) as instructed by the manufacturer. Briefly, 5uL of the Duo Transfection Reagent was added to 100uL of Opti-MEM (Gibco 31985062) medium and incubated at room temperature for 5 minutes. In a separate 100uL of Opti-MEM medium, 0.1pmol of BRCA2 expression plasmid and siRNA were mixed. The two mixtures were combined and incubated at room temperature for an additional 20 minutes. The combined mixture was added dropwise to cells for a total of 2mL of Opti-MEM in each well and a final siRNA concentration of 100nM. Six hours after transfection, the Opti-MEM medium was replaced with complete medium. Following 24 hours after transfection of oligonucleotides, cells were seeded at a density of 1000 cells per well of a 6-well plate in media containing either vehicle control (DMSO) or the respective concentration of Olaparib (LC Laboratories O-9201). Cells were incubated at 37°C for 10 days.

U2OS cells (1.5 × 10^5^) were transfected with either non-targeting control (Horizon Discovery, D-001206-14-05) or BRCA2 (Horizon Discovery) siRNAs at a final concentration of 100 nM using DharmaFECT 1 transfection reagent (Horizon Discovery, T-2001-02). Twenty-four hours following siRNA transfection, cells were transfected with 1µg of either WT or exon 11 deleted BRCA2 expression plasmids using Lipofectamine 2000 (Thermo Scientific, 11668019). Cells were plated (1,500 cells) on 6-well tissue dishes 24 h post-transfection, treated with olaparib and allowed to form colonies in presence of olaparib for 14 days.

For both HEK293T and U2OS cells, colonies were washed with 1× PBS followed by methanol fixation and staining with crystal violet staining solution (25% methanol + 0.5% crystal violet). Total number of colonies formed were normalized with the initial number of cells seeded and percent survival was calculated with respect to untreated control of individual experimental conditions. Graphs were generated using GraphPad Prism, version 10.2.3. p values were determined by Two-way ANOVA with Dunnett’s multiple comparisons test.

#### Immunofluorescence

HEK293T cells were co-transfected with BRCA2 siRNA and either an HA-tagged BRCA2 full-length or HA-tagged BRCA2 exon 11 deletion (BRCA2-ΔEx11) expression construct directly in Millicell EZ 8-well slides (Merck Millipore, 1,000 cells per well), as described previously. At 24 hours post-transfection, cells were treated with 5 μM Olaparib for 24 hours to induce DNA double-strand breaks. Cells were then washed with PBS, fixed with 4% paraformaldehyde for 15 min, and permeabilized with 0.5% Triton X-100 in PBS for 5 min. Samples were blocked for 40 min with blocking buffer (2% BSA, 0.2% Triton X-100, 1% goat serum in PBS). Primary antibodies diluted in blocking buffer were incubated overnight at 4°C: mouse monoclonal anti-HA (ab18181, Abcam; 1:2000) and rabbit monoclonal anti-RAD51 (ab133534, Abcam; 1:250). After three washes with PBS, cells were incubated with goat anti-mouse IgG Alexa Fluor 488 and goat anti-rabbit IgG Alexa Fluor 594 highly cross-adsorbed secondary antibodies (1:500 each; Thermo Fisher Scientific) for 1 hour at room temperature in the dark. Slides were mounted using ProLong Gold Antifade Reagent with DAPI (Invitrogen) and imaged on a Nikon Eclipse Ti2 microscope using a 100× objective. Contrast was adjusted uniformly across all images to improve foci visualization.

