## Supplementary Materials for "Landscape of reversion alterations in homologous recombination genes reveals evolutionary constraints"

**The PDF file includes:**

Supplementary Materials and Methods

Supplementary Figures S1 to S14

Supplementary Tables are provided in a separate .xlsx file.

Supplementary Materials and Methods

Oligonucleotides:

All siRNA were synthesized by Horizon Discovery (Waterbeach, United Kingdom)

BRCA2 siRNA sequence 1:	Sense: 5' C.G.U.U.U.A.A.A.G.C.C.A.A.G.A.A.U.A.A.U.U 3'

Antisense: 5' U.U.A.U.U.C.U.U.G.G.C.U.U.U.A.A.A.C.G.U.U 3'

BRCA2 siRNA sequence 2:	Sense: 5' C.C.A.C.A.A.A.A.G.C.A.G.A.A.G.A.U.U.A.U.U 3'

Antisense: 5' U.A.A.U.C.U.U.C.U.G.C.U.U.U.U.G.U.G.G.U.U 3'

WT BRCA2-HA pcDNA3.1(+): synthesized by GenScript (Nanjing, China)

Mutant (∆Exon 11) BRCA2-HA pcDNA3.1(+): synthesized by GenScript (Nanjing, China)

Small Molecules:

Olaparib: LC Laboratories O-9201

Transfection Reagents:

DharmaFECT Duo Transfection Reagent; Horizon Discovery T-2010-01

5X siRNA Buffer; Horizon Discovery B-002000-UB-100

Opti-MEM 1 Reduced Serum Medium; Gibco 31985062

Antibodies:

Anti-BRCA2: Millipore Sigma OP95

Anti-HA: Cell Signaling 3724

Anti-GAPDH: Cell Signaling 2118

Figure S1

Pan-cancer TBx and LBx deduplicated samples undergoing CGP between 2012 and 2024

*n=609,464*

TBx samples with one or more PVs in a gene in which REV were observed (*BRCA1, BRCA2, PALB2, RAD51C, RAD51D, BARD1, BRIP1,* or *CDK12)*

*n=75,417*

TBx samples with REV

*n=458*

LBx samples with one or more PVs in a gene in which REV were observed (*BRCA1, BRCA2, PALB2, RAD51C, RAD51D, BARD1, BRIP1,* or *CDK12)*

*n=20,363*

LBx samples with REV

*n=243*

Fig. S1. Consort diagram describing cross-sectional samples.

Figure S2

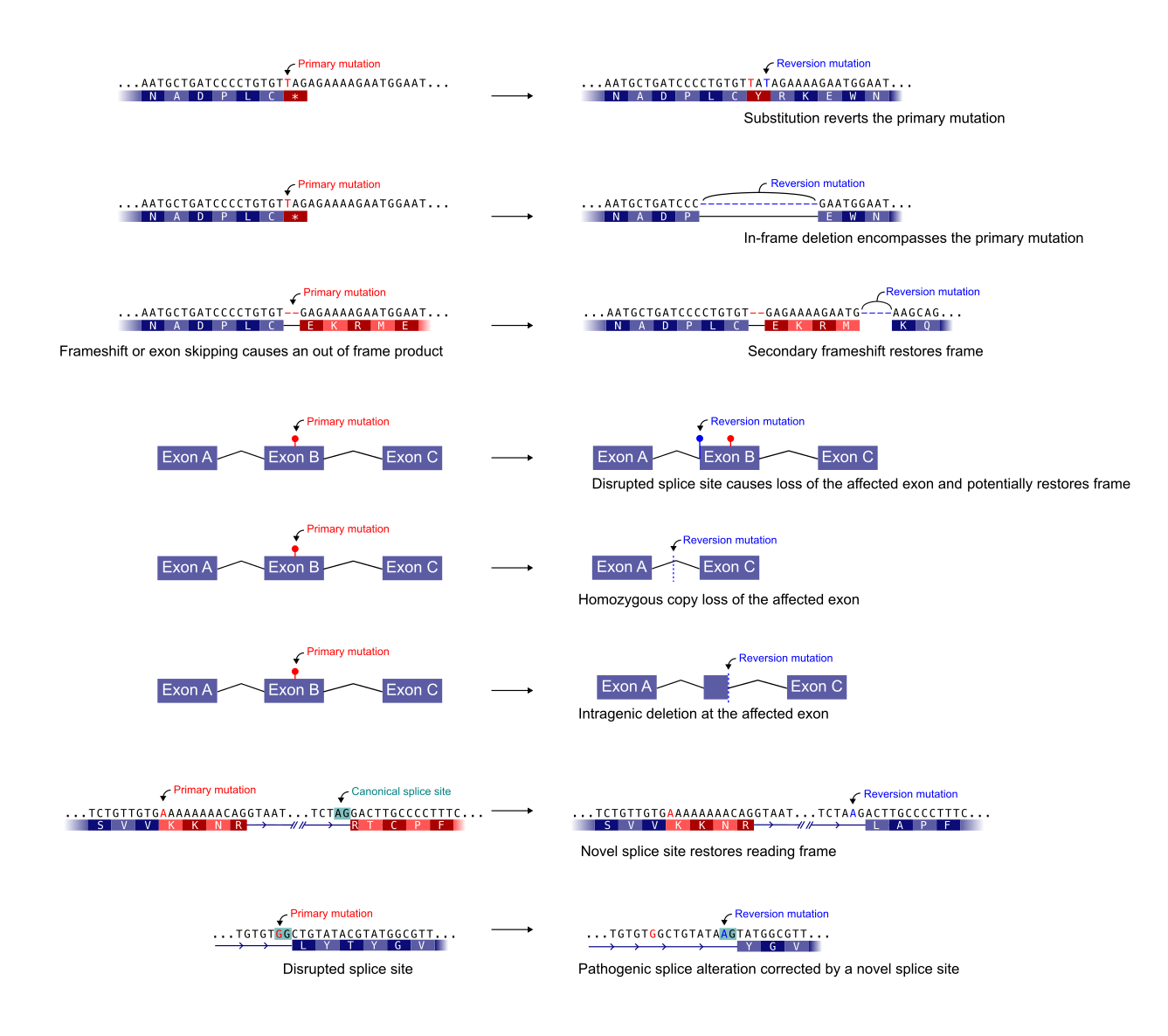

Fig. S2. Schema of REV mechanisms targeted for detection via the bespoke algorithm.

Figure S3

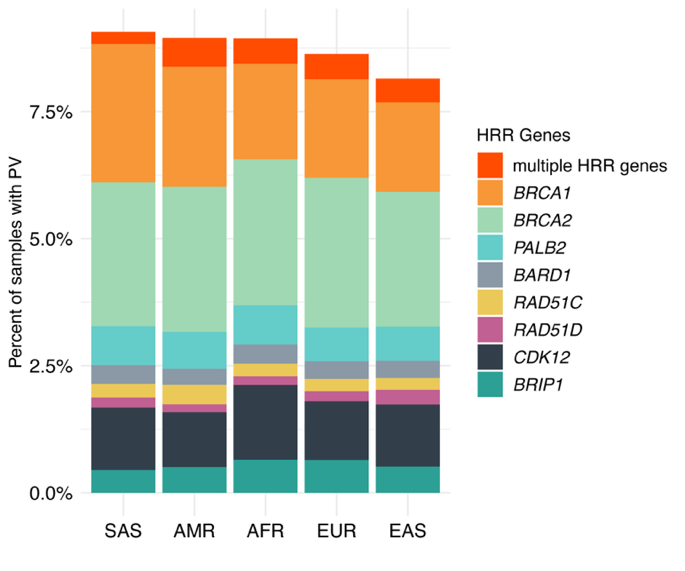

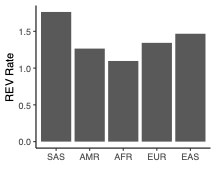

Fig. S3. Prevalence of pathogenic variants in REV8 genes across genomic ancestry superpopulations.

Figure S4

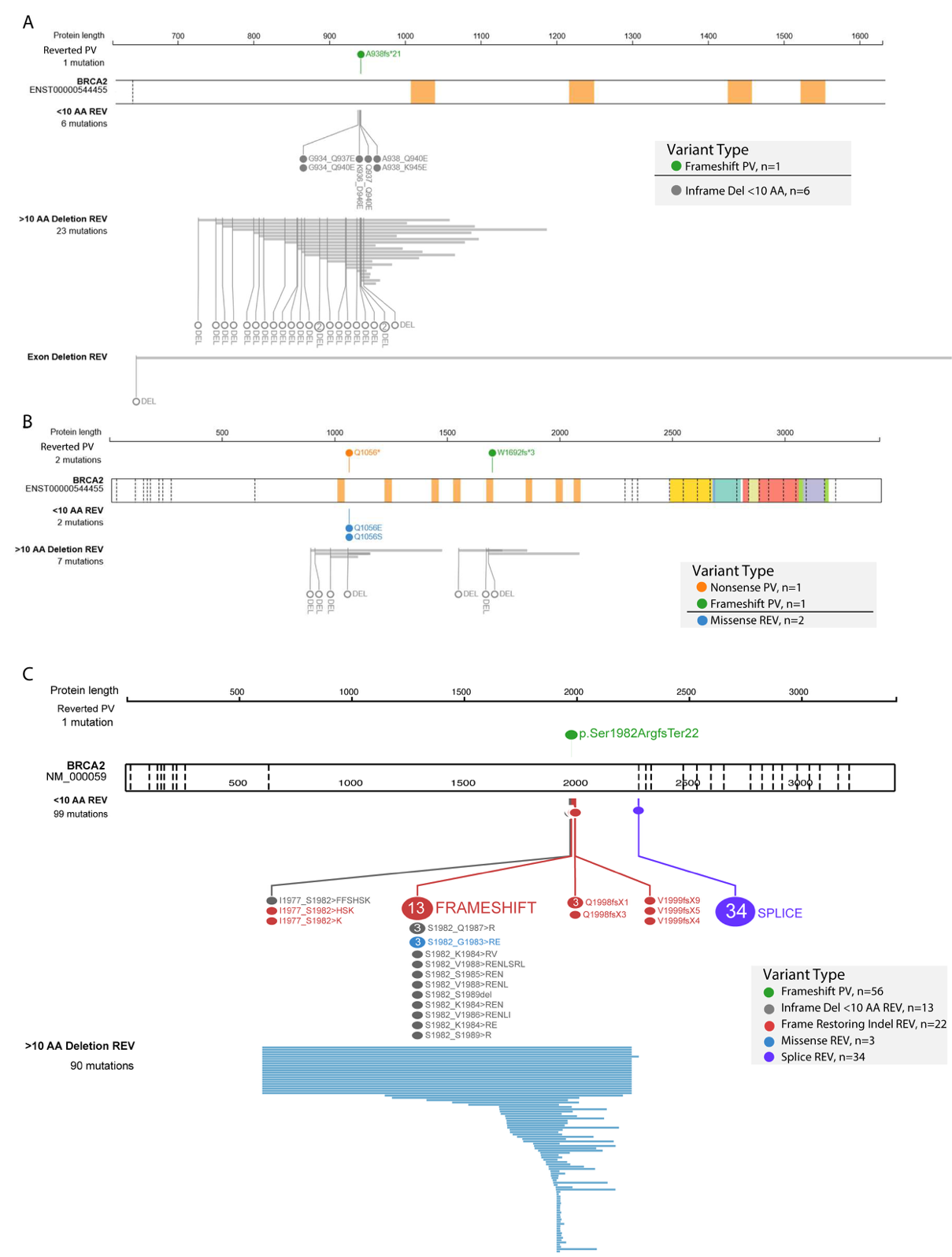

Fig. S4. Demonstrative examples of the evolutionary plasticity of REV. Numerous REV reverting a single PV in one LBx sample (A), multiple PV independently reverted (B), and cross-sample REV predicted to restore function to the Ashkenazi Jewish founder allele *BRCA2* p.Ser1982ArgfsTer22. The upper track contains REV <10 AA in length and the lower track depicts inframe deletion REV within spanning >10 AA in length.

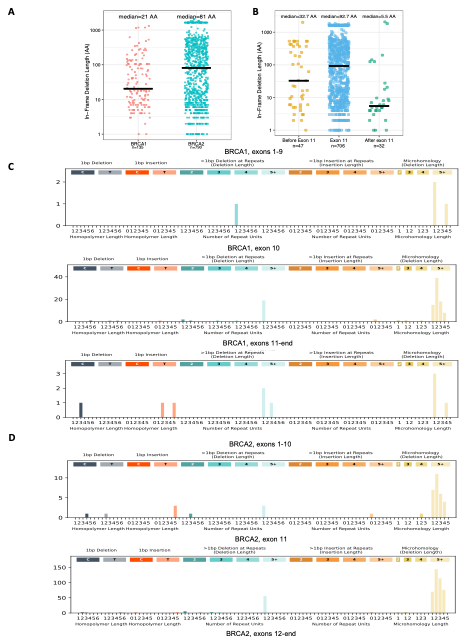
Figure S5

Fig. S5. Length and microhomology characteristics of indel REVs in *BRCA1* and *BRCA2*. (A) Length of in frame deletion REVs by gene. (B) Length of in-frame deletion REVs by position in both *BRCA1* and *BRCA2*. Indel signatures for mutations causing REVs by location in *BRCA1* (C) and *BRCA2* (D), including counts of mutations in three categories: 1bp insertion or deletion in the context of increasing homopolymer length (slate and fire), insertions or deletions larger than 1 bp in repetitive regions by length of the repeat (light blue/orange), and deletions that occur with microhomology by length of the microhomology region (yellow). Complex deletion-insertion REVs were excluded.

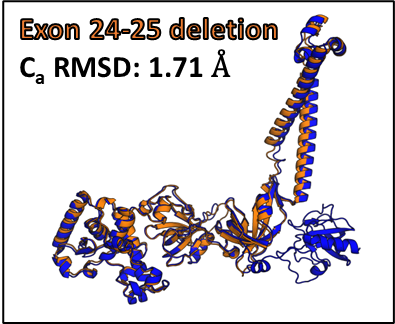

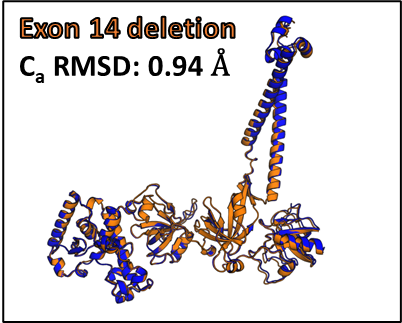

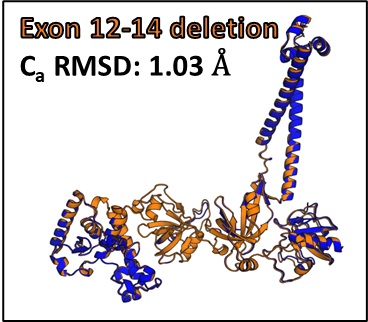

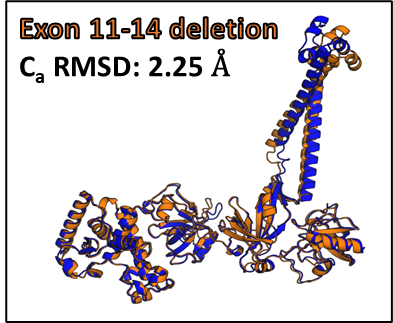

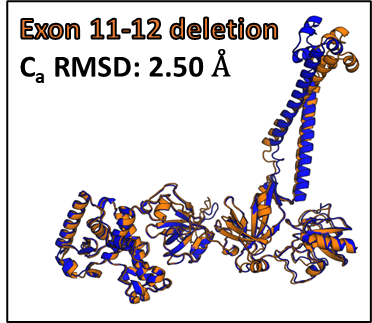

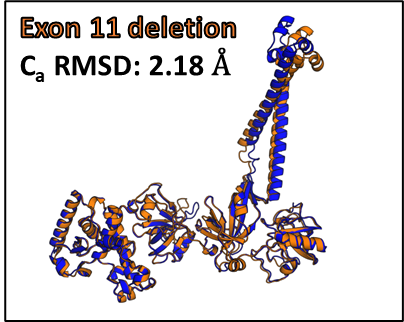

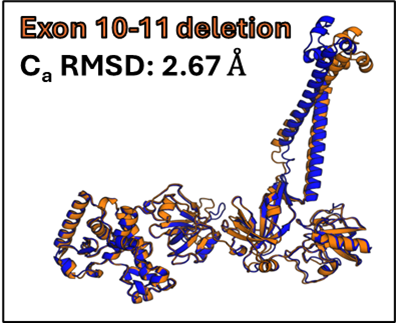

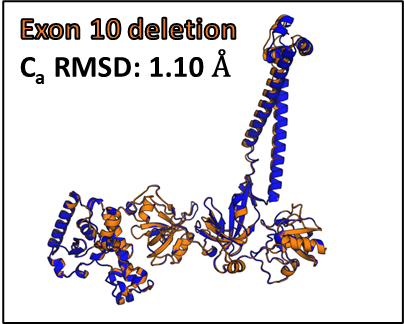

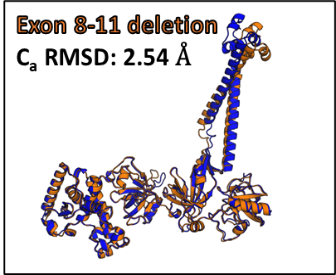

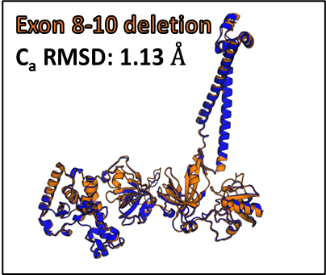

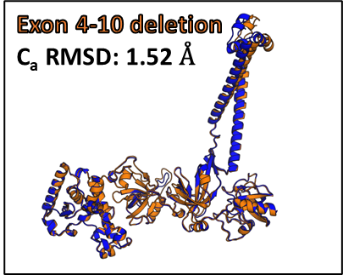

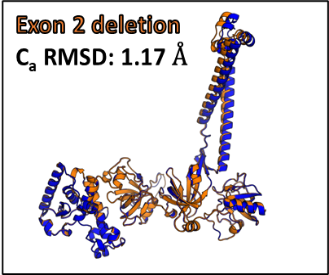

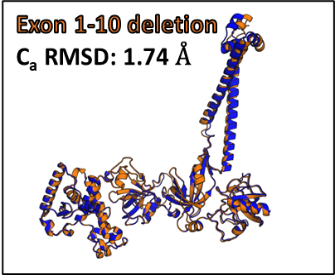
Figure S6

**BRCA2 WT**

**BRCA2 exon(s) deletion**

**C_a_ RMSD for exons 15-25**

Fig. S6. Top-ranked AlphaFold model for each BRCA2 mutant (shown in orange) and AlphaFold-generated wildtype BRCA2 model (shown in blue). The RMSD values between wildtype BRCA2 and the mutants were calculated in MSL1 using alpha carbon atoms from exons 15-25. The RMSD was calculated using exons 15-23 for the exon 24-25 deletion mutant.

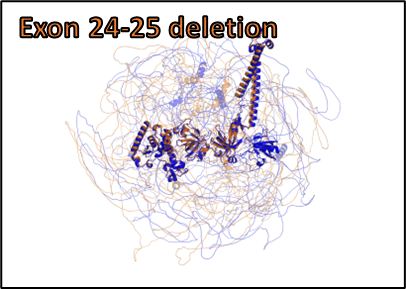

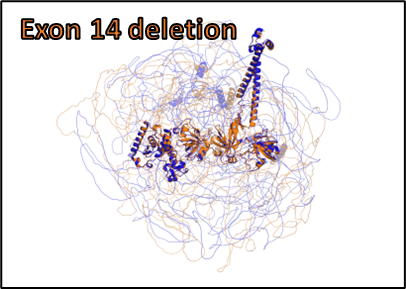

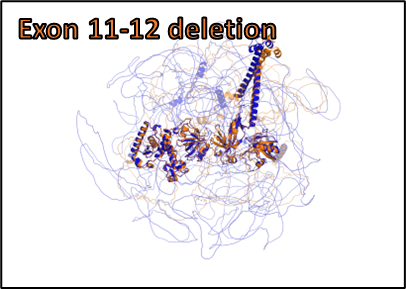

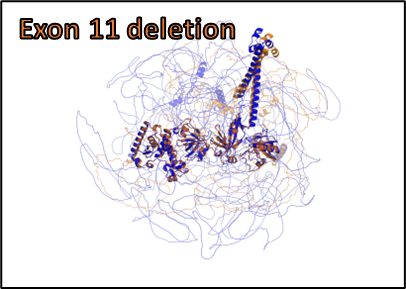

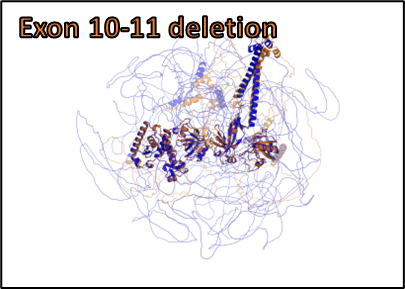

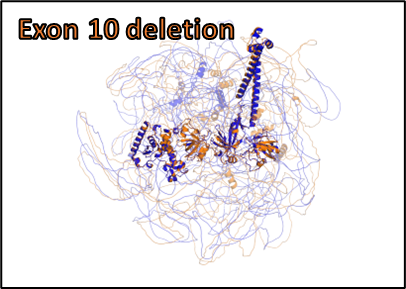

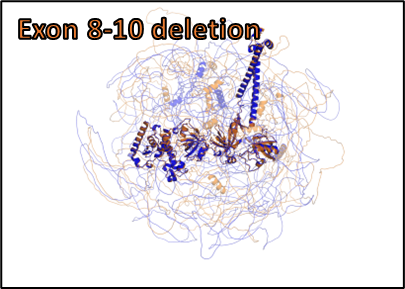

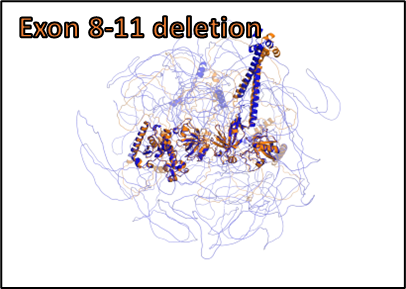

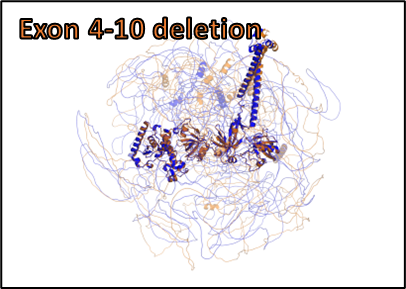

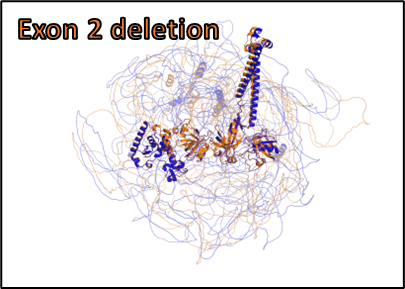

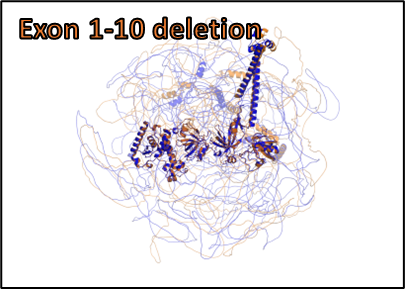
Figure S7

**BRCA2 WT**

**BRCA2 exon(s) deletion**

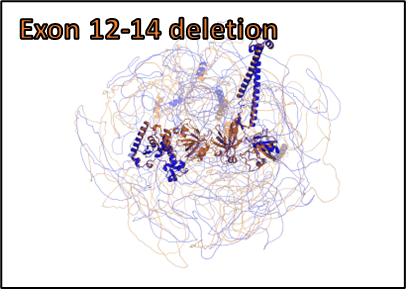

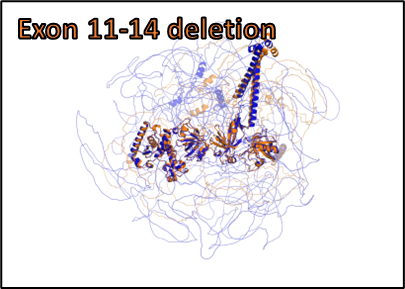

Fig. S7. Top-ranked AlphaFold model for each BRCA2 mutant (shown in orange) and AlphaFold-generated wildtype BRCA2 model (shown in blue). Exons 15-25 (15-23 for the exon 24-25 deletion mutant) are highlighted while all other exons are transparent. The wildtype and mutants were aligned using exons 15- 25 in PyMOL.

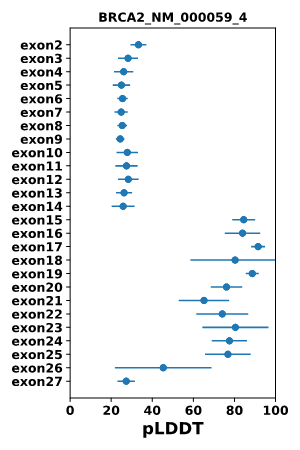

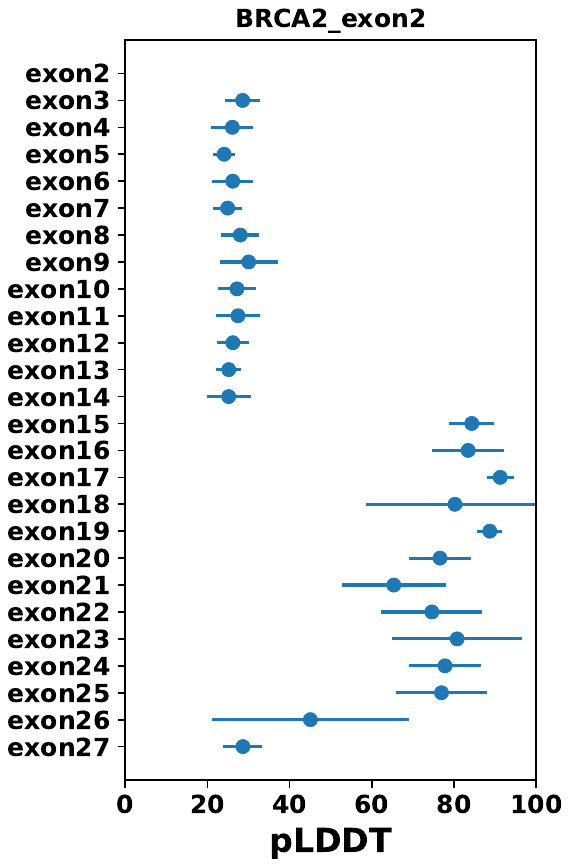

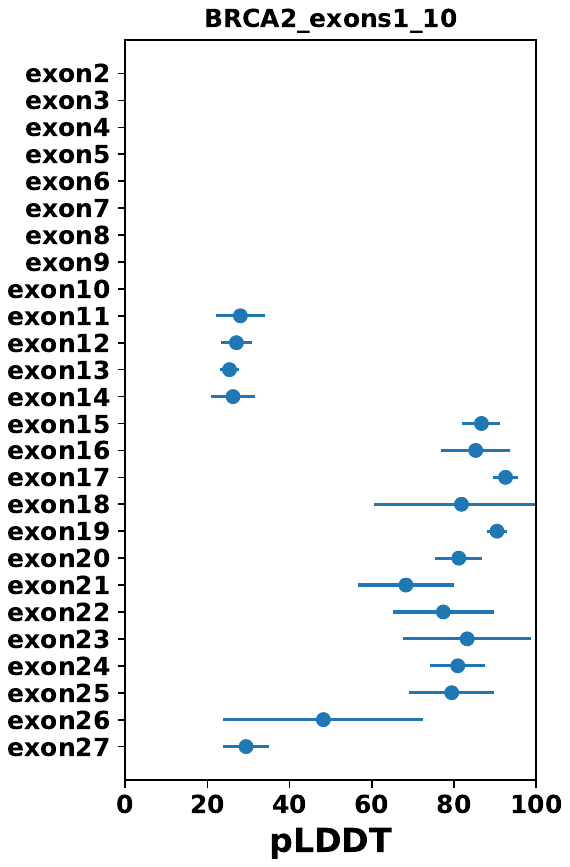

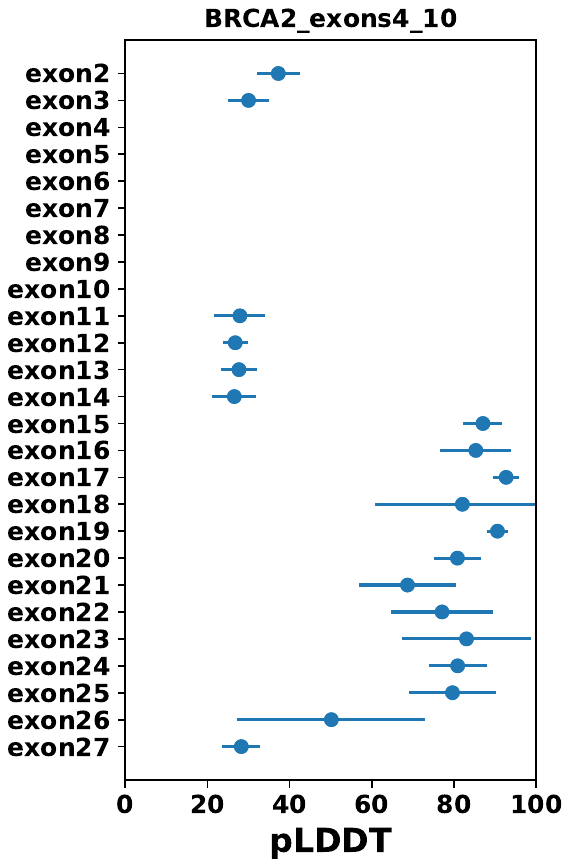
Figure S8

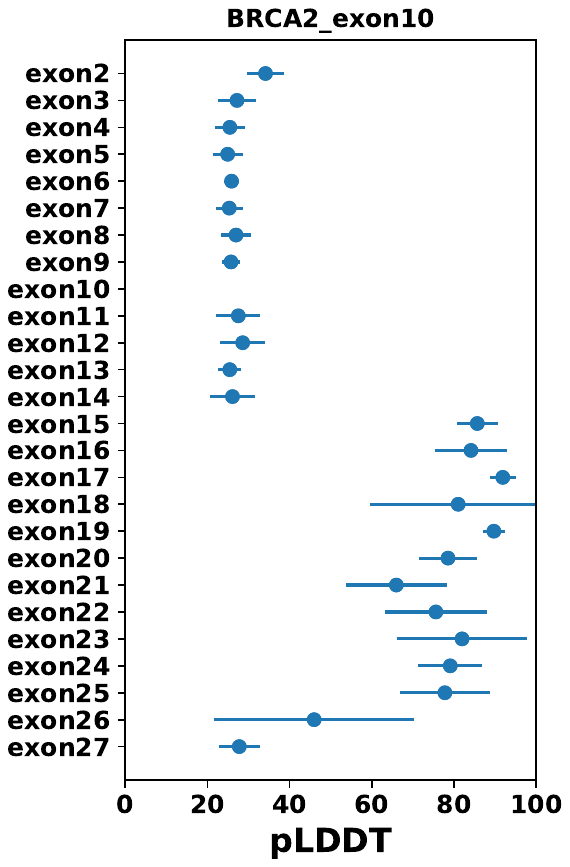

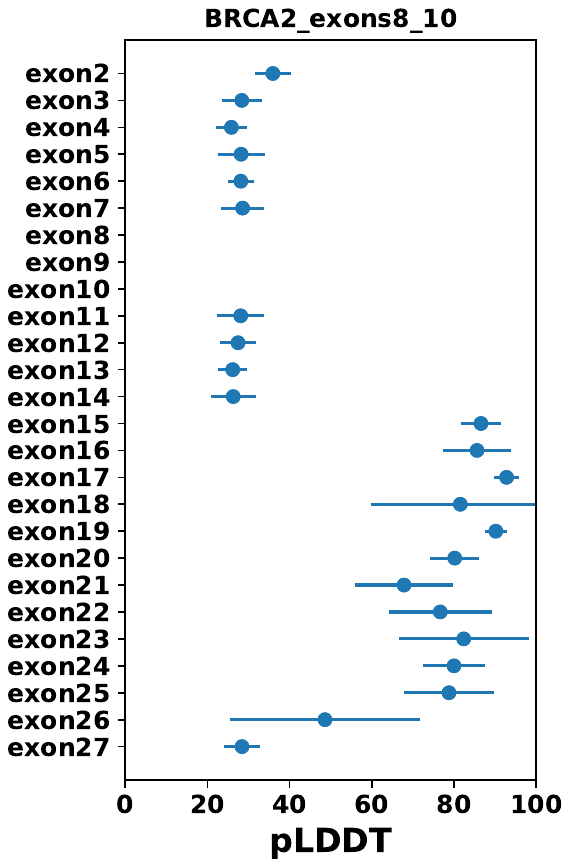

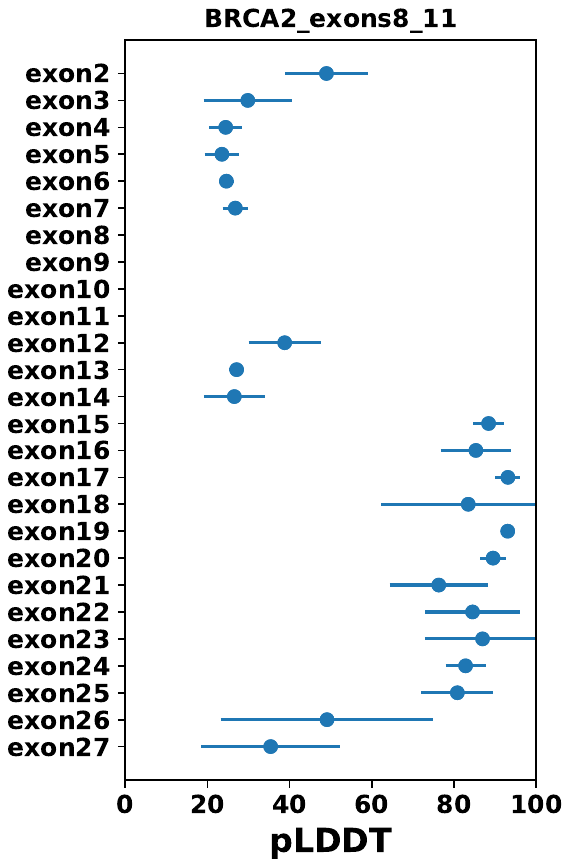

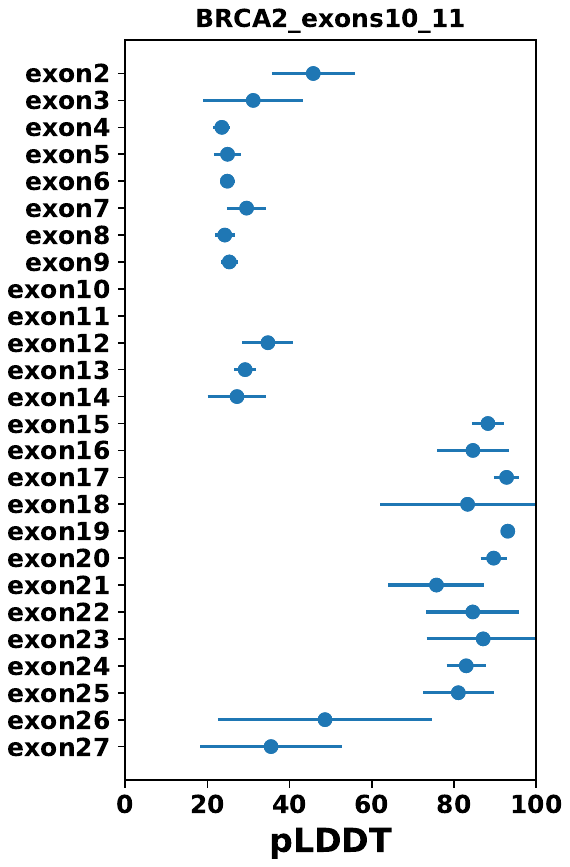

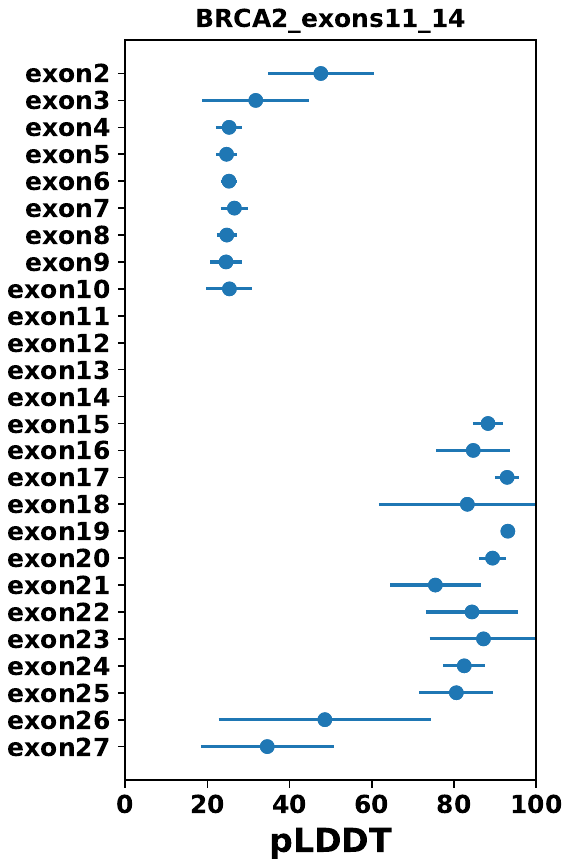

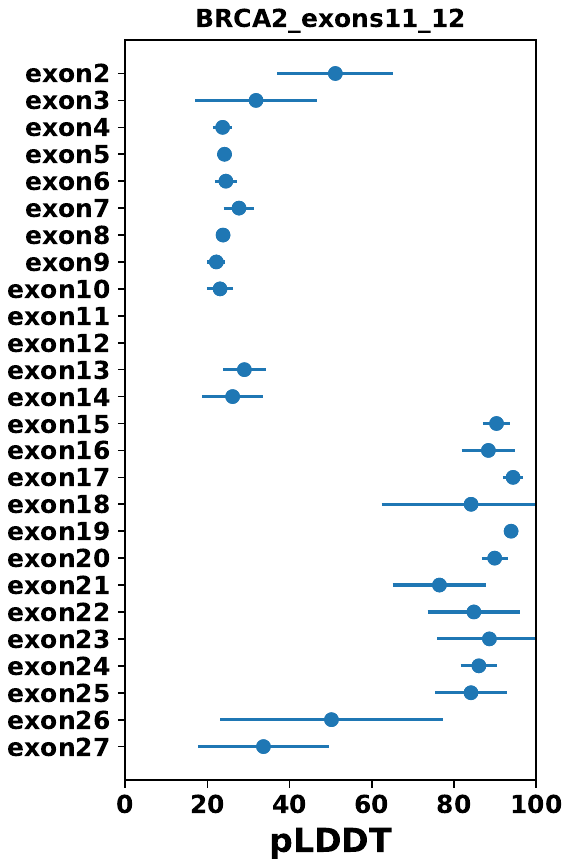

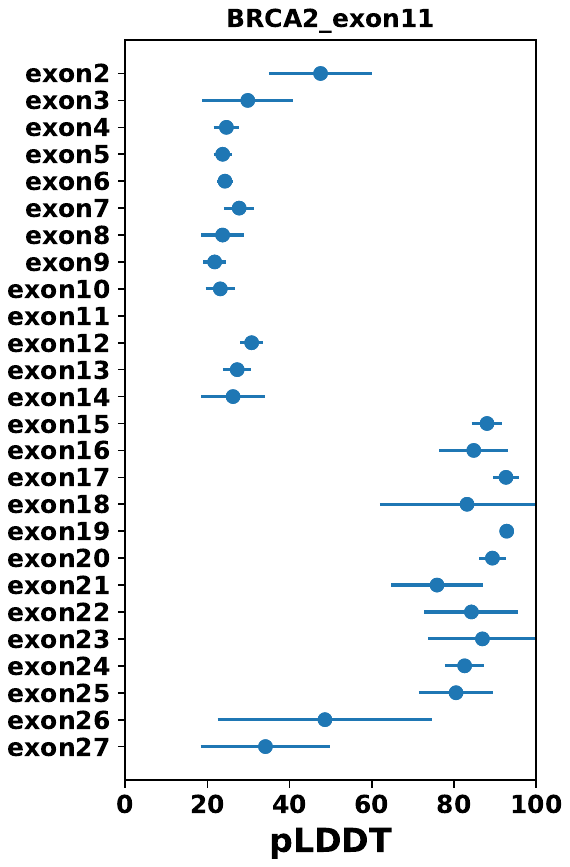

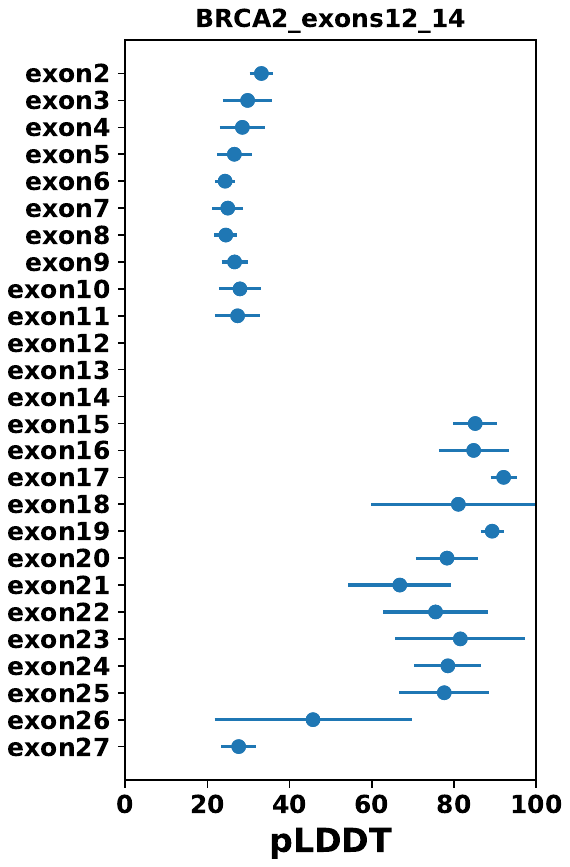

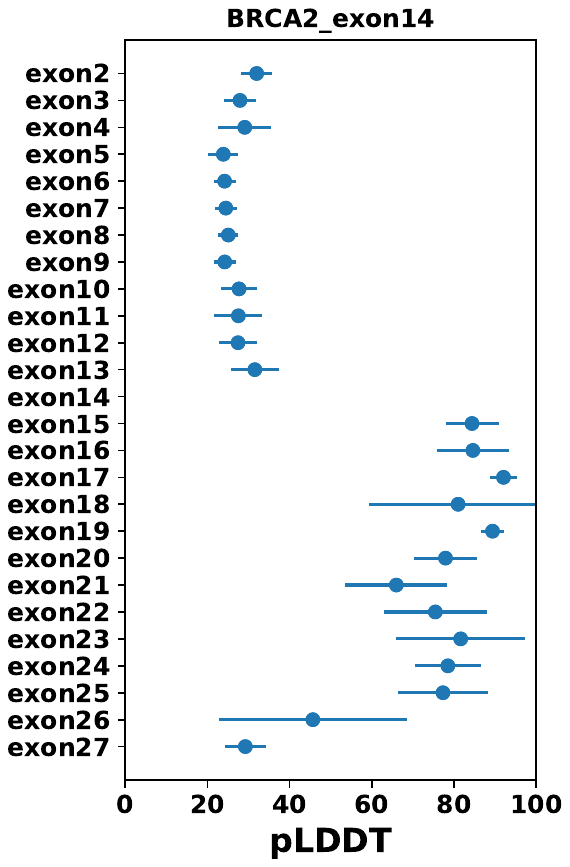

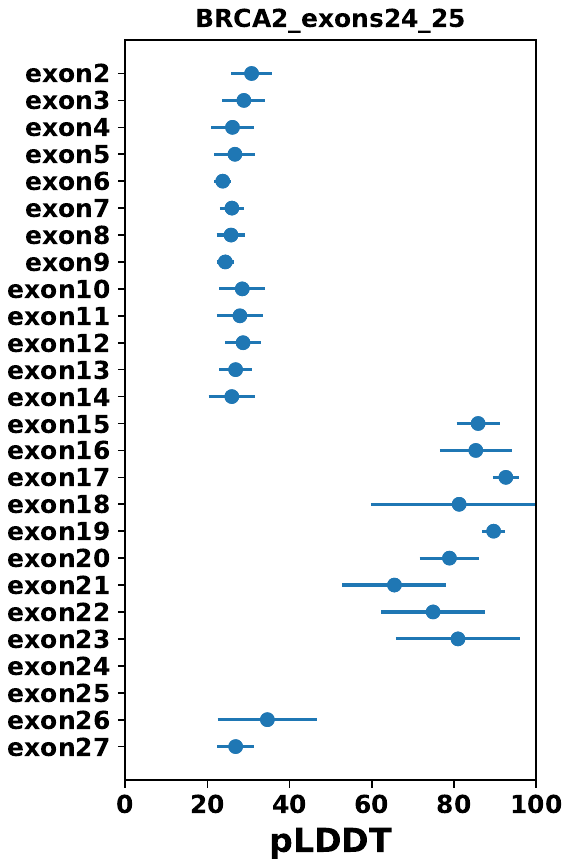

Fig. S8. Average pLDDT values per BRCA2 exon, with error bars showing 1 standard deviation

Figure S9

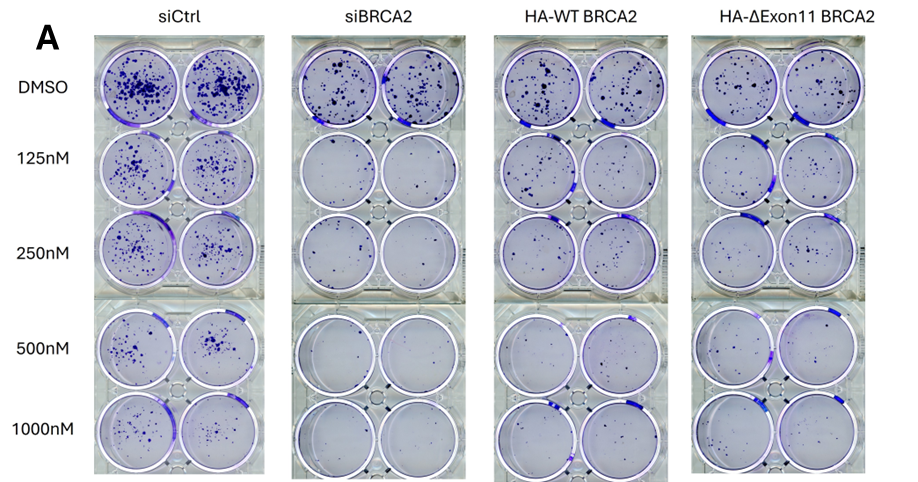

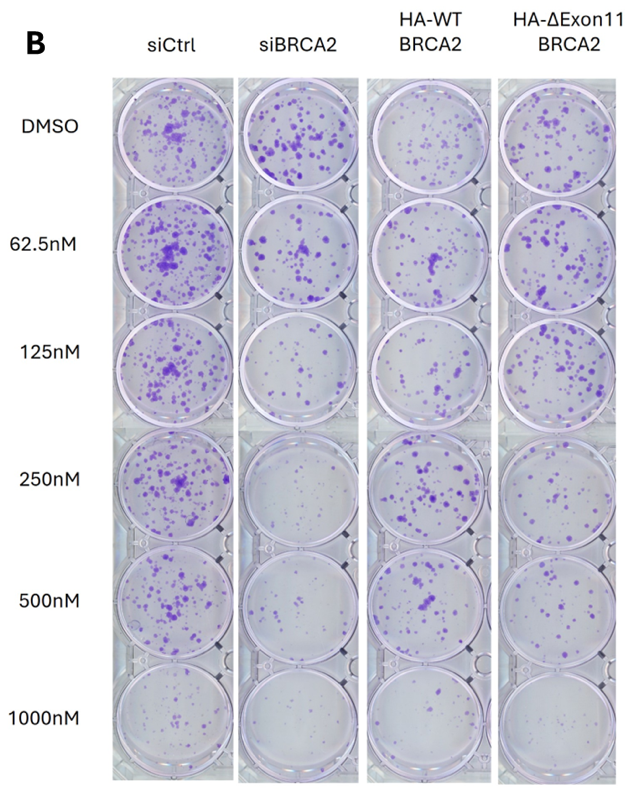

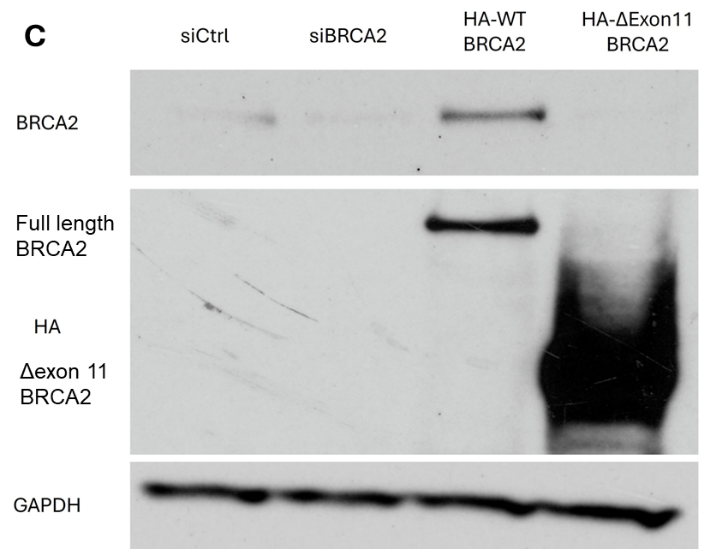

Fig. S9. Response to olaparib comparing siControl, siBRCA2, siBRCA2 plus HA - WT BRCA2, siBRCA2 plus HA - Δexon 11 BRCA2. A) Example of colony formation studies in HEK 293T cells. B) Example of colony formation studies in U2OS cells. C) Western blot demonstrating expression of full length BRCA2 and Δexon 11 BRCA2.

Figure S10

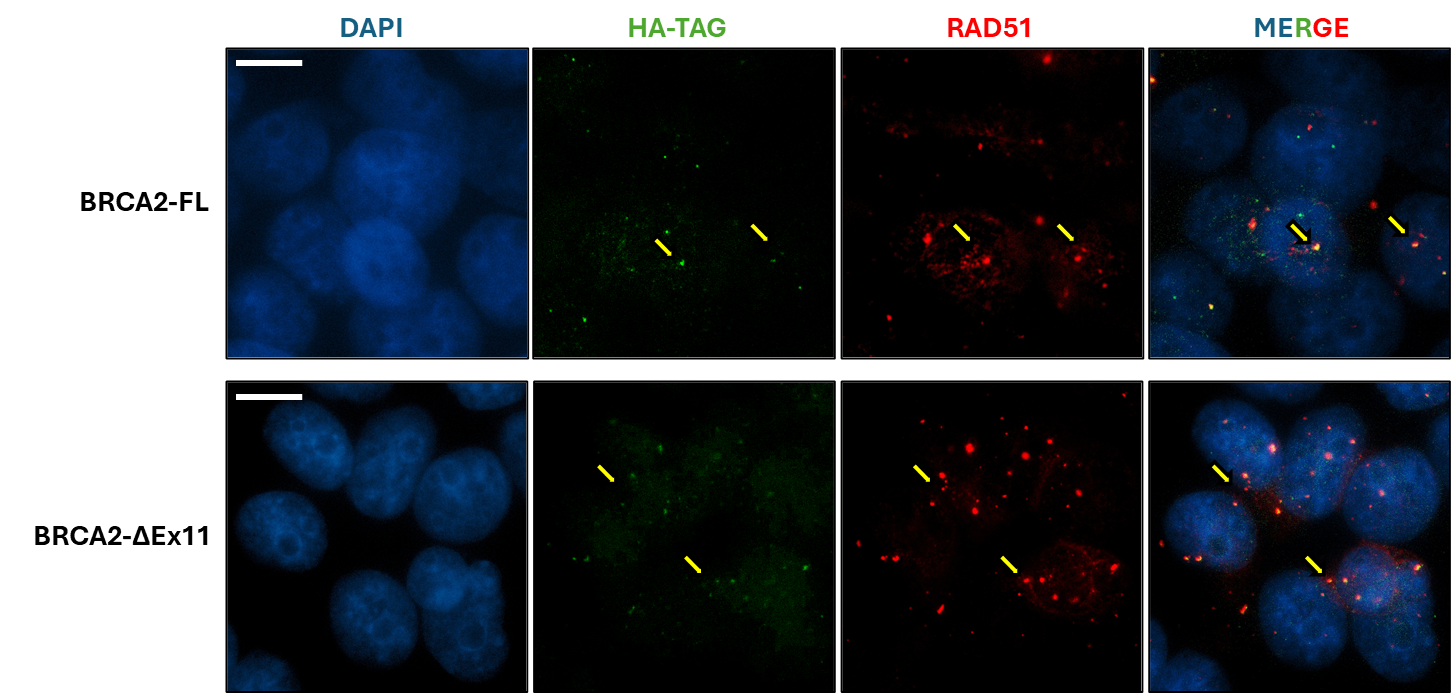

Fig. S10. BRCA2 full-length and BRCA2-ΔEx11 proteins co-localize with RAD51 foci following DNA damage. Immunofluorescence images of HEK293T cells co-transfected with BRCA2 siRNAs and either an HA-tagged BRCA2 full-length (BRCA2-FL) or HA-tagged BRCA2 exon 11 deletion (BRCA2-ΔEx11) rescue construct. Cells were treated with 5 μM Olaparib for 24 hours prior to fixation to induce DNA double-strand breaks. Channels shown: DAPI (blue), HA-tag (green, Alexa Fluor 488), RAD51 (red, Alexa Fluor 594), and merged. Yellow arrows indicate sites of co-localization between HA-tagged BRCA2 constructs and RAD51 foci. Representative images are shown. Images were acquired on a Nikon Eclipse Ti2 microscope with a 100x objective. Contrast was adjusted uniformly across all images to aid visualization of foci. Scale bar, 10 μm.

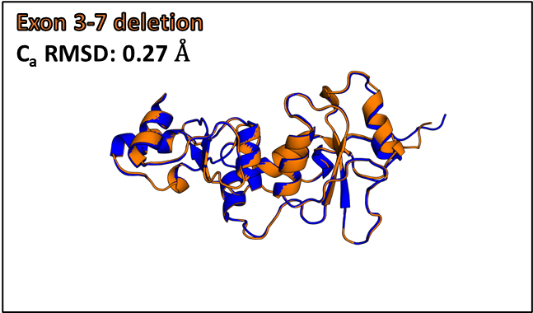

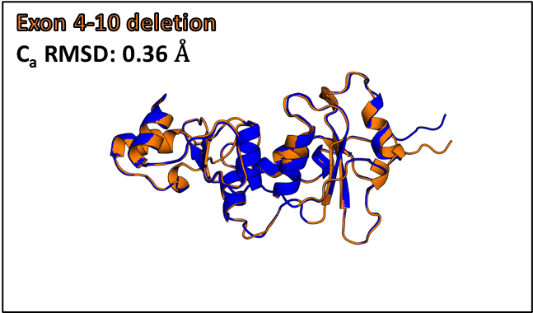

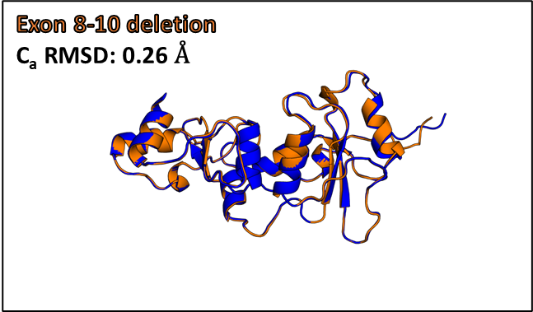

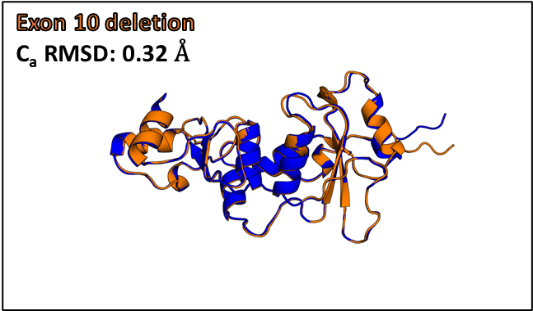

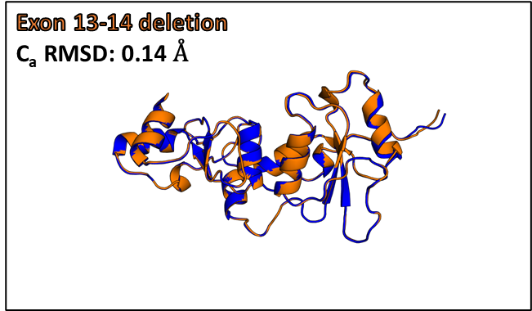

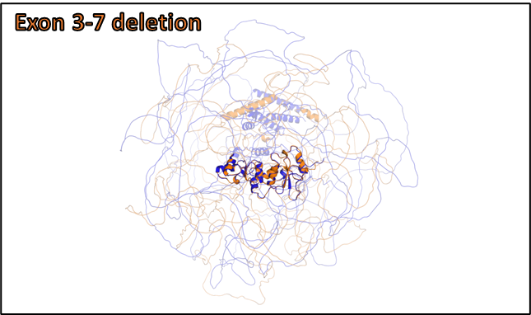

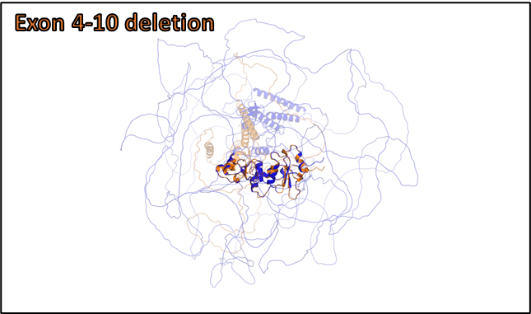

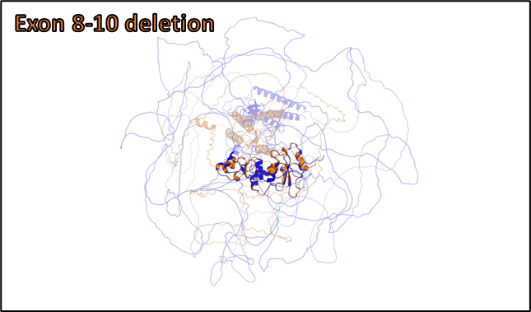

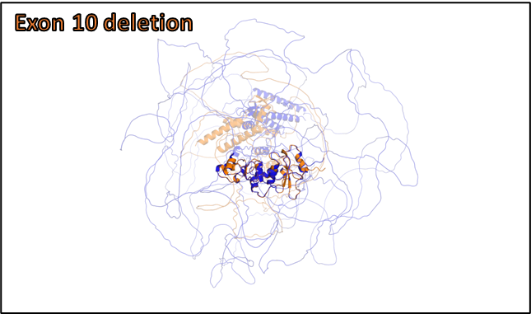

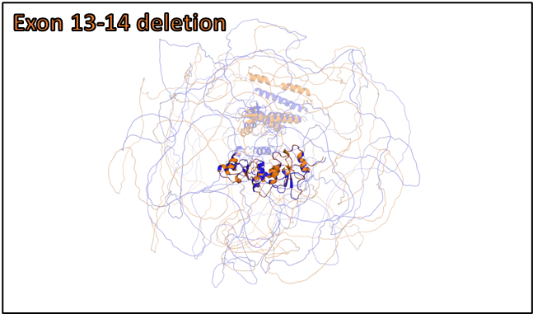
Figure S11

Fig. S11. Top - The top-ranked AlphaFold model for each BRCA1 mutant is shown in orange, the AlphaFold-generated wildtype BRCA1 model is shown in blue. The RMSD values between wildtype BRCA1 and the mutants were calculated in MSL using alpha carbon atoms from exons 16-23. Bottom - The top-ranked AlphaFold model for each BRCA1 mutant is shown in orange, the AlphaFold-generated wildtype BRCA1 model is shown in blue. Exons 16-23 are highlighted while all other exons are transparent. The wildtype and mutants were aligned using exons 16-23 in PyMOL.

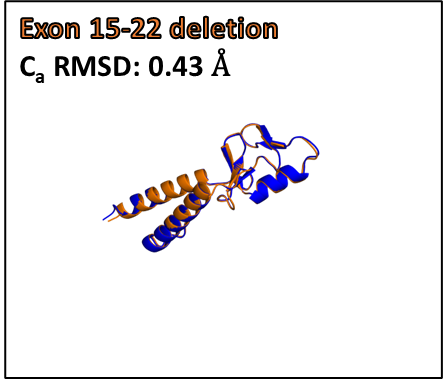

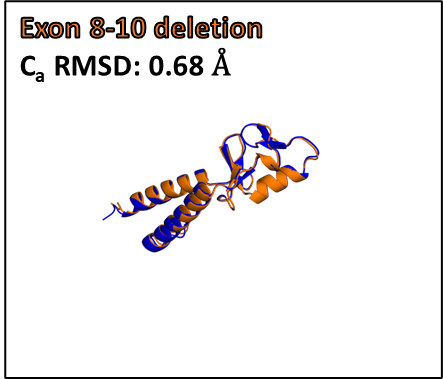

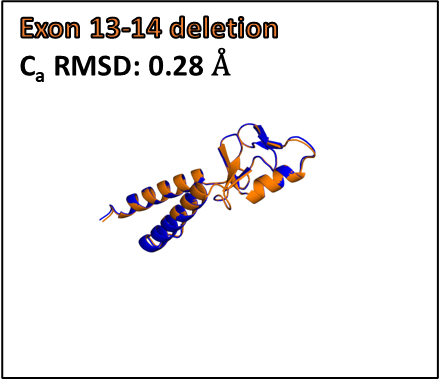

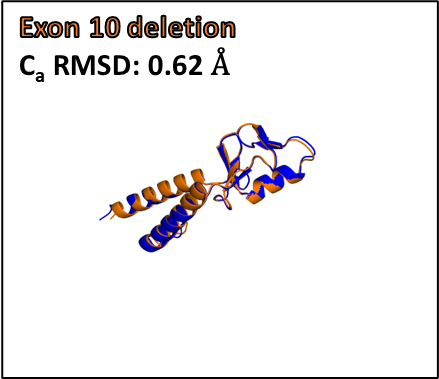
Figure S12

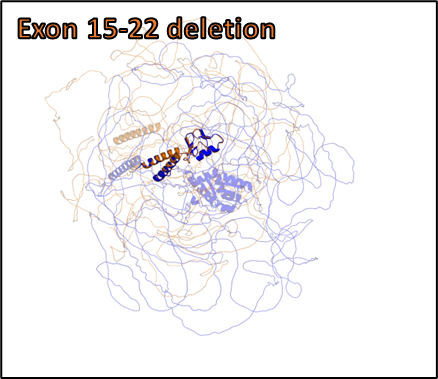

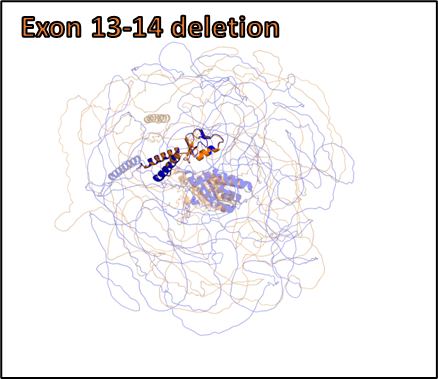

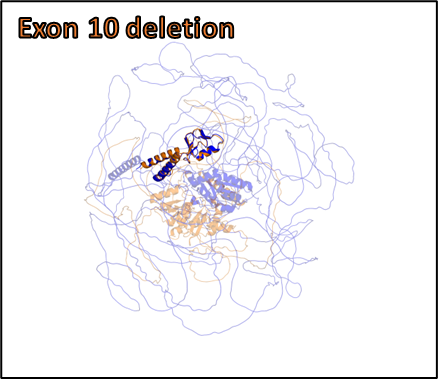

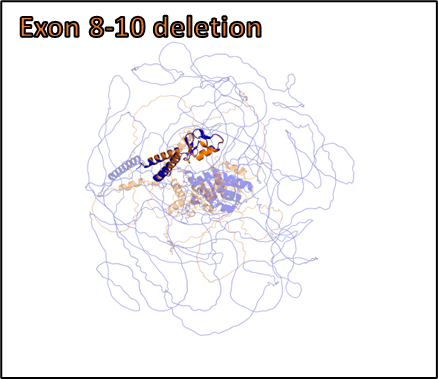

Fig. S12: Top - The wildtype and mutants were aligned using exons 2-5 in PyMOL. The RMSD values between wildtype BRCA1 and the mutants were calculated in MSL using alpha carbon atoms from exons 2-5. Bottom - The top-ranked AlphaFold model for each BRCA1 mutant is shown in orange, the AlphaFold-generated wildtype BRCA1 model is shown in blue. Exons 2-5 are highlighted.

Figure S1
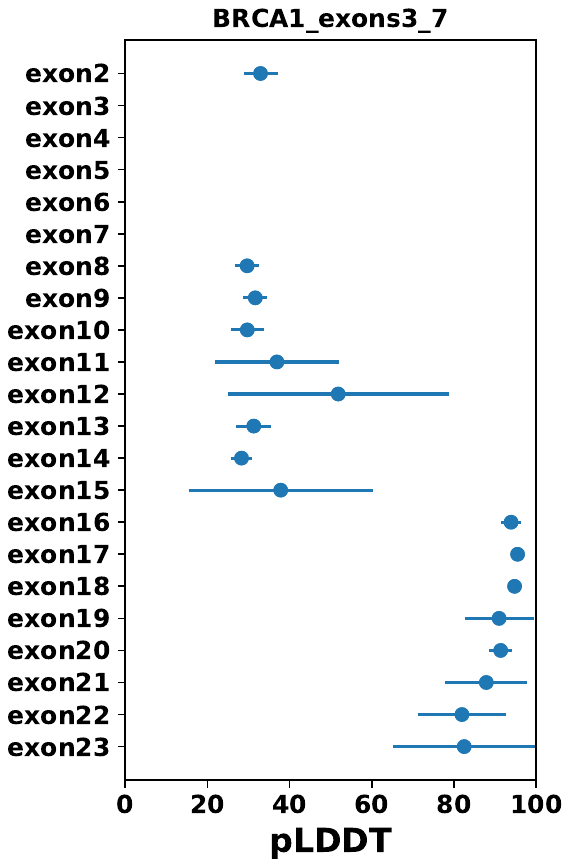

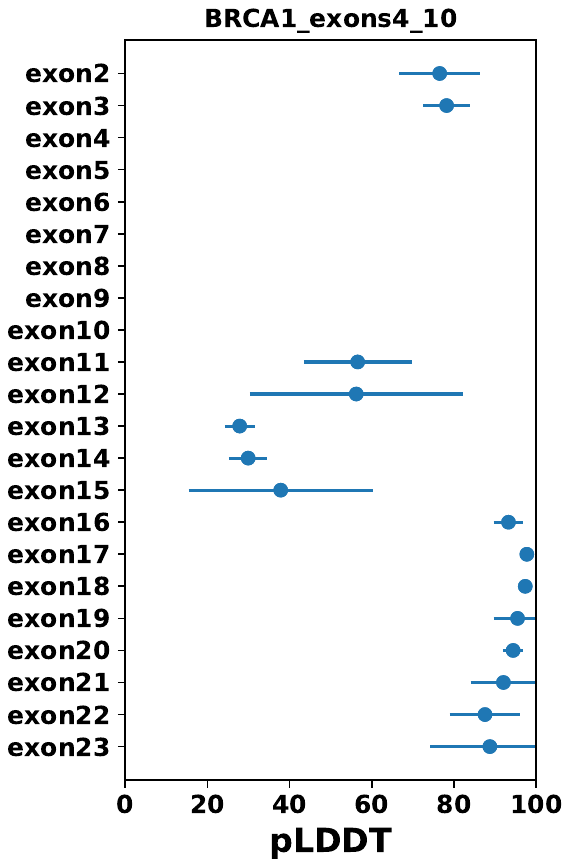

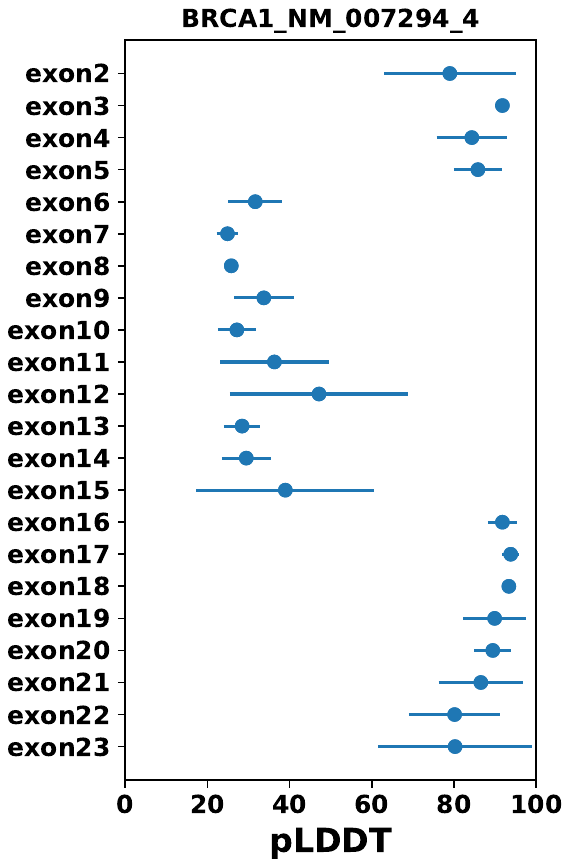
3

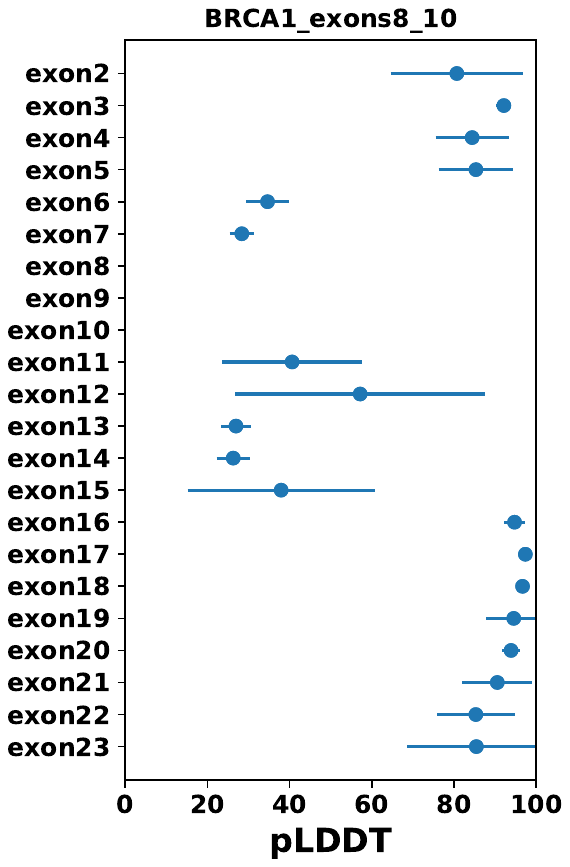

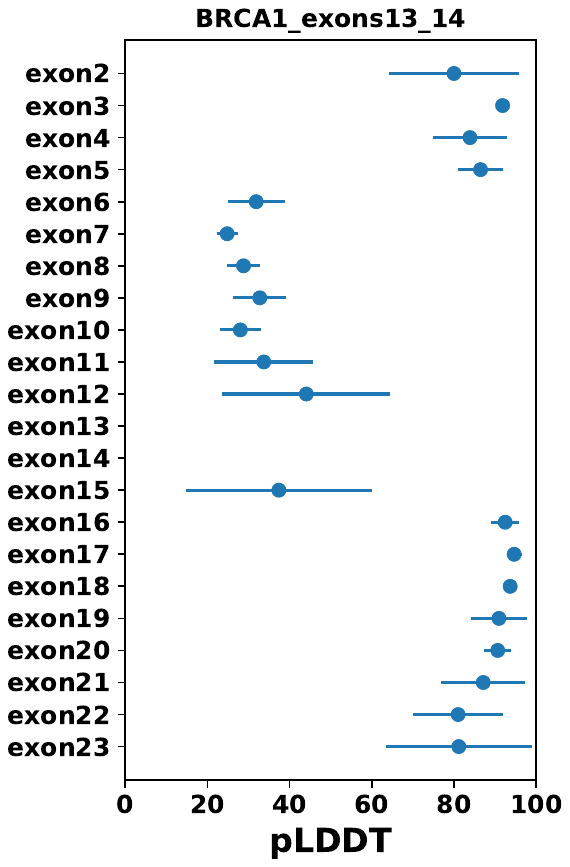

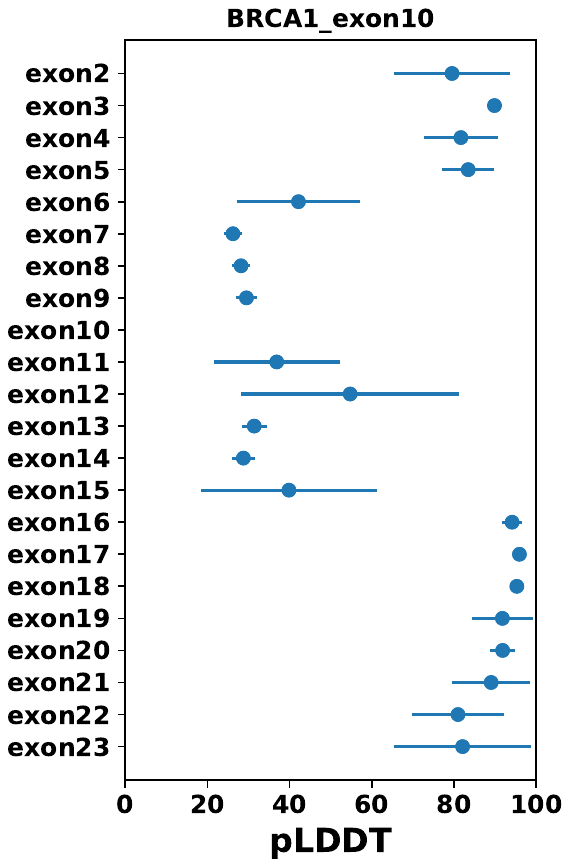

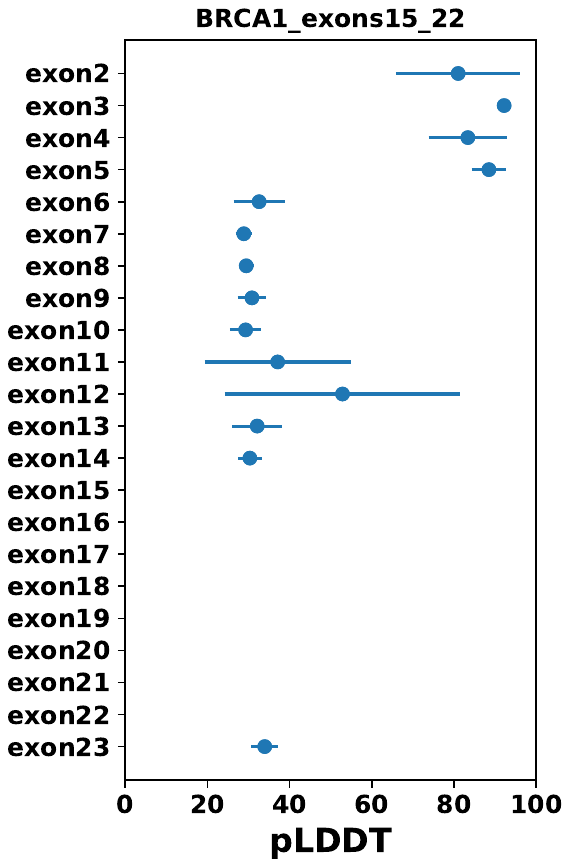

**Fig. S13. Average pLDDT values per exon for BRCA1, with error bars showing 1 standard deviation.**

Figure S14

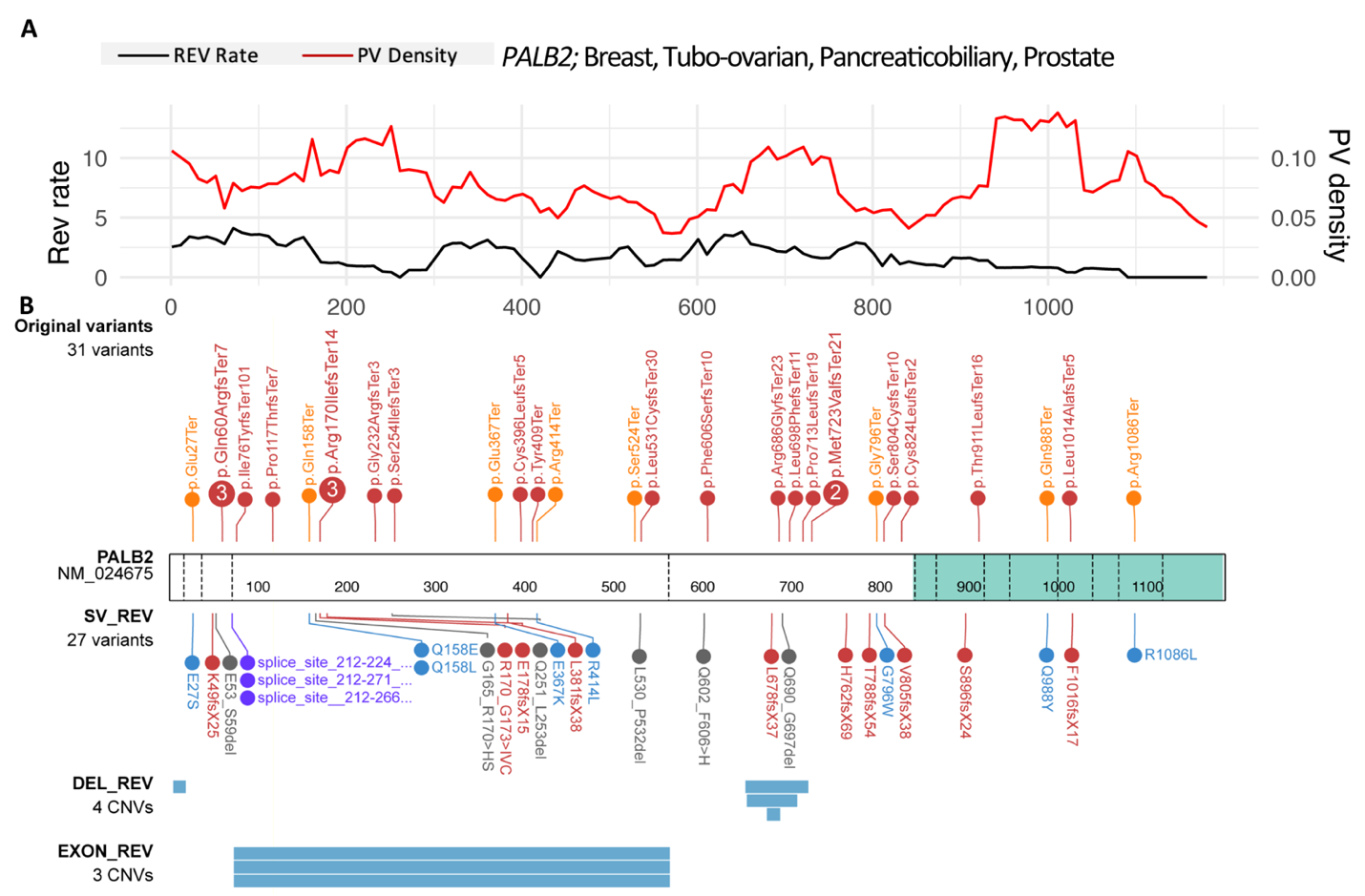

Fig. S14: Characteristics of REVs involving *PALB2*. A) REV rate and PV density within 100-AA sliding windows by PV AA position for breast, tubo-ovarian, prostate, and pancreatic cancers. B) Reverted PV type and position (track above the gene model) and REV mechanism and position (tracks below the gene model), with the upper track containing REVs <10 AA in length, the middle track depicting inframe deletion REVs within exons and spanning >10 AA in length, and the bottom track displaying exon-level or larger deletion REVs.
